# Epithelial Morphogenesis Driven by Cell-Matrix vs. Cell-Cell Adhesion

**DOI:** 10.1101/2020.06.24.165795

**Authors:** Shaohe Wang, Kazue Matsumoto, Kenneth M. Yamada

## Abstract

Many embryonic organs undergo epithelial morphogenesis to form tree-like hierarchical structures. However, it remains unclear what drives the budding and branching of stratified epithelia, such as in embryonic salivary gland and pancreas. Here, we performed live-organ imaging of mouse embryonic salivary glands at single-cell resolution to reveal that budding morphogenesis is driven by expansion and folding of a distinct epithelial surface cell sheet characterized by strong cell-matrix adhesions and weak cell-cell adhesions. Profiling of single-cell transcriptomes of this epithelium revealed spatial patterns of transcription underlying these cell adhesion differences. We then synthetically reconstituted budding morphogenesis by experimentally suppressing E-cadherin expression and inducing basement membrane formation in 3D spheroid cultures of engineered cells, which required β1 integrin-mediated cell-matrix adhesion for successful budding. Thus, stratified epithelial budding, the key first step of branching morphogenesis, is driven by an overall combination of strong cell-matrix adhesion and weak cell-cell adhesion by peripheral epithelial cells.

## INTRODUCTION

Branching morphogenesis is widely used by epithelial organs to maximize their functional surface area (Wang et al., 2017). All branching organs have a core epithelium encased by a layer of basement membrane, which is surrounded by a loosely condensed group of mesenchymal cells. Among various roles, the mesenchyme secretes growth factors critical for epithelial growth and morphogenesis (Affolter et al., 2009; Costantini and Kopan, 2010; Patel et al., 2006; Shih et al., 2013; Wang et al., 2017). However, when suitable growth factors and extracellular matrix are provided, the epithelium of many organs can branch without the mesenchyme (Ewald et al., 2008; Nogawa and Ito, 1995; Nogawa and Takahashi, 1991), indicating the core capacity for branching is intrinsic to the epithelium.

Branching epithelia can be single-layered with a lumen or stratified without a lumen. Branching of a single-layered epithelium involves buckling of the epithelial sheet (Nelson, 2016). The buckling of single-layered lung epithelium can be guided by external sculpting forces from airway smooth muscle cells (Goodwin et al., 2019; Kim et al., 2015), although other cell types are likely involved in vivo (Young et al., 2020). In stratified epithelia, however, the concept of buckling cannot be easily applied to account for branching morphogenesis due to the apparent lack of a sheet-like structure.

Embryonic salivary gland and pancreas are classical examples of stratified epithelia that undergo branching morphogenesis, which comprises distinct phases of budding and ductal morphogenesis (Shih et al., 2013; Steinberg et al., 2005; Wang et al., 2017). During budding morphogenesis, numerous epithelial buds arise from repeated clefting of a single initial epithelial bud, whereas ductal morphogenesis generates the epithelial tubular structures connecting terminal end buds together. Budding morphogenesis is characterized by extensive dynamics of epithelial cells and the basement membrane matrix (Harunaga et al., 2014; Larsen et al., 2006; Shih et al., 2016), but it remains unclear how numerous epithelial buds arise from the interplay of cell and matrix dynamics or other mechanisms.

Here, we use volumetric live-organ imaging to follow individual cells within virtually the entire mouse embryonic salivary gland during branching morphogenesis. We find that surface-localized epithelial cells form an integral layer with the basement membrane, which together expands and folds inward to drive budding morphogenesis. We use numerical modeling and experimental perturbations to corroborate a model that a combination of weak cell-cell adhesion and strong cell-matrix adhesion of peripheral epithelial cells drives the expansion and folding of the surface epithelial sheet. Furthermore, we employ single-cell RNA sequencing and single-molecule RNA FISH (fluorescence in situ hybridization) to reveal distinct transcriptional features of these surface epithelial bud cells. Importantly, we demonstrate successful reconstitution of budding morphogenesis by experimentally reducing E-cadherin expression and inducing basement membrane formation in 3D spheroid cultures of engineered epithelial cells that normally do not form buds. Our results reveal a fundamental self-organizing mechanism based on preferential cell-matrix adhesion vs. cell-cell adhesion that can explain how stratified epithelia undergo budding morphogenesis.

## RESULTS

### Clefting in salivary glands is caused by uniform expansion and inward folding of the surface cell sheet

To visualize cellular mechanisms of stratified epithelial branching, we developed live-organ imaging strategies using two-photon microscopy that enabled us to image nearly the entire 3D volume of transgenic mouse embryonic salivary glands at high spatiotemporal resolution (**Fig. S1A-B; Video S1**). 3D cell tracking revealed extensive cell motility throughout the developing gland with cell migration rates increasing near the periphery of the branching epithelial buds as previously described (Hsu et al., 2013; Larsen et al., 2006) (**Fig. S1C**).

Next, we evaluated whether cells exchange freely between the outer epithelial layer and gland interior during morphogenesis, or whether branching salivary glands are composed of distinct interior and surface cell populations. To do so, we photoconverted patches of cells near the epithelial surface in transgenic salivary glands expressing KikGR, a photoconvertible fluorescent protein emitting green or red fluorescence before or after conversion (Hsu et al., 2013; Tsutsui et al., 2005). Most photoconverted peripheral epithelial cells moved rapidly along the tissue surface while maintaining intimate contact with the basement membrane (**Fig. 1A; Video S2**), suggesting tight adherence of these cells to the encasing basement membrane. Furthermore, we used an epithelial RFP reporter (Krt14p::RFP) that exhibited elevated expression in peripheral versus interior epithelial cells (**Fig. S1D**) to enable automated rendering of the epithelial surface (**Fig. 1B-C**). We analyzed cell movements at the epithelial surface (located within 15 μm of the gland surface at any point within the tracked time window), which revealed that the movements of most epithelial cells near the surface remain confined to the surface of the developing tissue (**Figs. 1D-E, S1C; Video S3**). During new bud formation by clefting, the peripherally enriched Krt14p::RFP reporter clearly delineated a distinct surface cell sheet, whose expansion and folding seemed to underlie clefting (**Video S4**).

**Figure 1.**
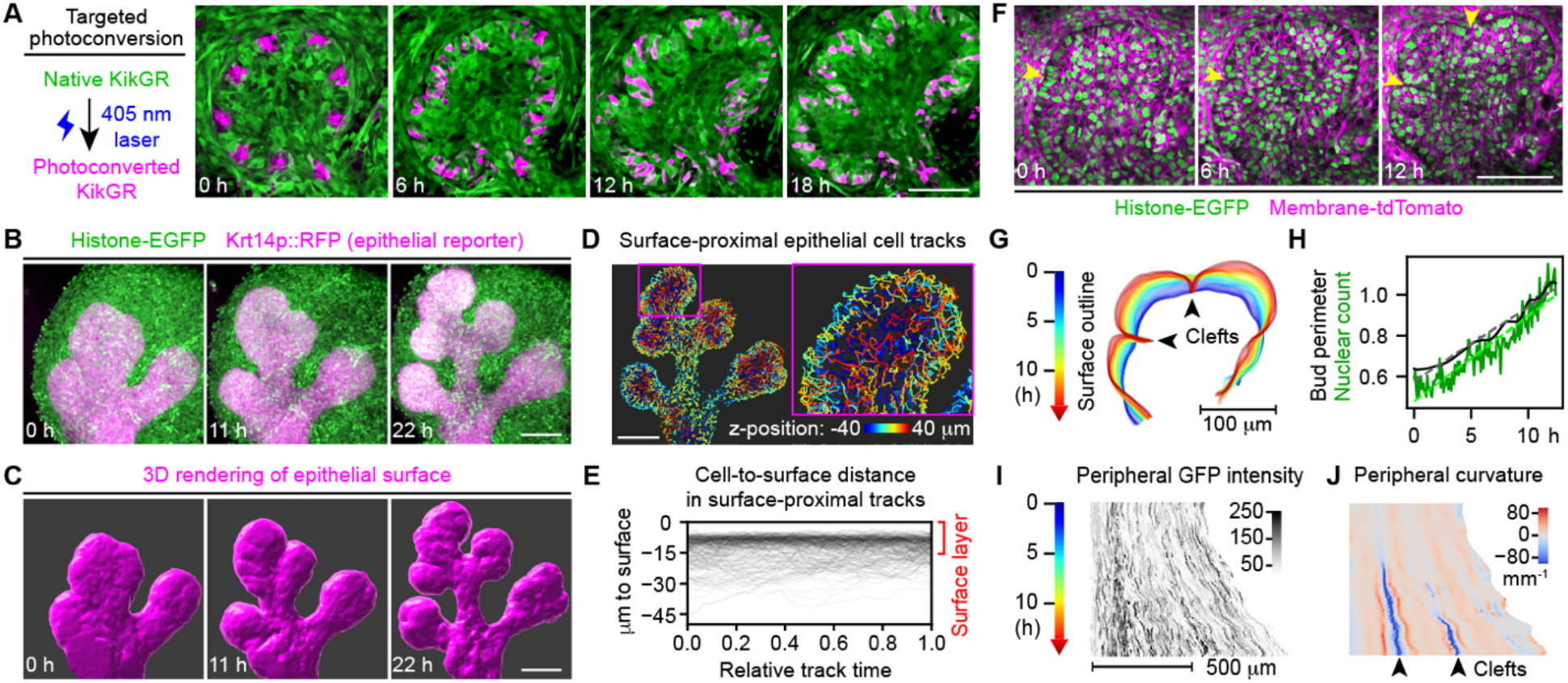
Clefting in salivary glands is caused by uniform expansion and inward folding of the surface cell sheet. (**A**) Left: schematic of KikGR photoconversion; Right: snapshot confocal images showing the middle slice of a branching epithelial bud in an E13 mouse salivary gland expressing KikGR. (**B**) Time-lapse two-photon microscopy images showing the maximum intensity projection of an E12.5 transgenic mouse salivary gland. (**C**) 3D rendering of epithelial surface using Krt14p::RFP at time points matching the images in (B). (**D**) Surface-proximal epithelial cell tracks (tracking nuclear Histone-EGFP) color-coded by their z-position at 20-22 hours of the time-lapse sequence. Only epithelial cell tracks whose closest distance to the surface was ≤ 15 μm are shown. (**E**) Plot of the cell nucleus-to-surface distance versus time for 250 randomly selected 3-10 hour-long surface-proximal tracks. (**F**) Time-lapse two-photon microscopy images showing the middle slice of an E13 transgenic mouse salivary gland. (**G**) Outlines of the epithelial surface at the middle two-photon image slice over 12.5 hours at 5-min intervals. Blue to red, 0 to 12.5 hours. (**H**) Plot of the bud perimeter and nuclear count along the surface cell layer at the middle slice over time. Dashed lines indicate fitted linear models. (**I** and **J**) Heatmaps of GFP intensity (I) and the curvature (J) along the surface epithelial cell layer at the middle slice over time. Arrowheads in (F, G, J) indicate clefts. Scale bars, 100 μm.

Next, we determined whether new surface cells are added uniformly around the epithelial surface or locally at the cleft in order to distinguish between clefting as a systemic or local process. We traced nuclear histone-EGFP intensities of peripheral epithelial cells over time and computed local peripheral curvature to track surface deformation (**Figs. 1F-J, S1D**). Local expansion to form a cleft would predict an abrupt change in slope angles of temporal nuclear traces at cleft sites (**Fig. S1E**). However, the observed changes of slope angles were gradual, and surface expansion rates near clefts were indistinguishable from other locations, suggesting that clefting is a systemic activity (**Figs. 1I-J, S1H-J**). Moreover, increasing peripheral nuclear counts over time closely matched expansion of the bud perimeter, indicating constant peripheral cell density (**Figs. 1H, S1F-G, K**).

Taken together, we conclude that clefting in salivary glands is caused by uniform expansion and inward folding of the surface epithelial cell sheet.

### Expansion of the surface cell sheet is driven by subsurface cell division and reinsertion as new surface cells

We next determined the origin of new epithelial surface cells. The distinct boundary of Krt14p::RFP expression levels between peripheral and interior epithelial cells hinted that new surface cells arise primarily from proliferation of preexisting surface cells (**Video S4**). However, none of the 289 surface cells whose division was monitored divided within the surface layer to produce two surface daughter cells (Type III; **Fig. 2A**). Instead, 92.4% of the surface cells moved to a subsurface level to complete cell divisions that produced two daughter cells in the gland interior (Type I; **Figs. 2A, S2A**), and the remaining 7.6% divided in an orientation perpendicular to the surface to generate one surface daughter cell and one interior daughter cell (Type II; **Figs. 2A, S2B**). Importantly, all surface cell-derived interior daughter cells eventually returned to the surface by reinserting between surface cells, causing a delayed surface expansion (**Figs. 2B, S2C-D; Video S5**). Most cells returned to the surface within 4 hours, but some required >12 hours (**Fig. 2B**). The location of daughter cell reinsertion could be distant from the parental surface cell. In one example, a daughter cell meandered under a forming cleft into a neighboring bud to reach the bottom surface distant from the parental cell location (**Fig. S2E**). Overall, the reinsertion sites were uniformly distributed around the epithelial surface (red dots in **Fig. S2D**), revealing the cellular basis of uniform surface expansion.

**Figure 2.**
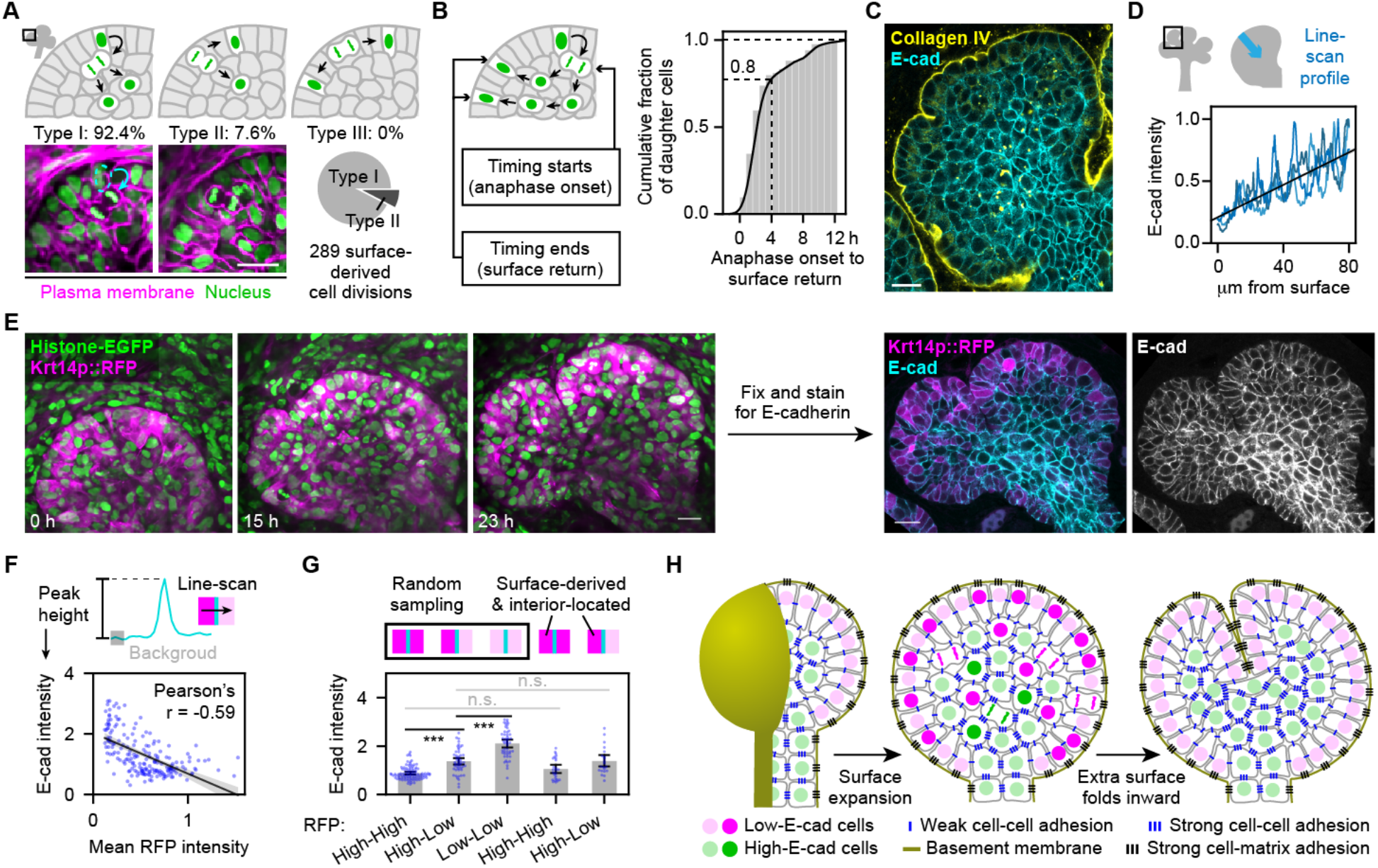
Expansion of the surface cell sheet is driven by subsurface cell division and reinsertion as new surface cells. (**A**) Top and lower left: schematics and time-lapse two-photon microscopy images of 3 types of surface-derived cell division; Lower right: pie chart showing proportions of the two observed types. (**B**) Schematic and cumulative distribution plot of time intervals from anaphase onset of mother cell division to returning of 84 daughter cells to the epithelial surface. (**C**) Confocal immunofluorescence image showing the middle slice of an epithelial bud from an E13.5 mouse salivary gland. (**D**) Schematic and plot of the surface-to-center line-scan profile of E-cadherin intensity. (**E**) Left: time-lapse two-photon microscopy images showing the middle slice of a branching epithelial bud in an E13 transgenic mouse salivary gland. Right: confocal immunofluorescence images showing the middle slice of the same epithelial bud at 23 h. (**F**) Top: schematic of how E-cadherin intensity was measured. Bottom: scatter plot of the E-cadherin intensity of an edge vs. the mean RFP intensity of its two adjacent cells. Black line is the linear regression with 95% confidence interval shown in gray shading. (**G**) Plot of E-cadherin intensity at indicated categories of cell-cell boundaries. Error bars, 95% confidence intervals. ***, Tukey test p<0.001. n.s., not significant. (**H**) Schematic model of clefting in a stratified epithelium. First, the surface layer expands by surface cell division and back-insertion. Second, the extra surface folds inward to produce clefting. Note that the two steps happen concurrently but are drawn separately for clarity. Scale bars, 20 μm.

What drives the robust surface return of surface-derived cells? Based on the lower E-cadherin expression level of peripheral epithelial cells compared to interior epithelial cells (Walker et al., 2008) (**Fig. 2C-D**), we hypothesized that differential cell-cell adhesion directed sorting out of low E-cadherin surface-derived cells from high E-cadherin interior cells (Steinberg, 1963).

To determine whether surface-derived cells maintained low E-cadherin expression when they were temporarily interior-located after cell division, we fixed transgenic salivary glands (Krt14p::RFP and Histone-EGFP) immediately after live imaging and immunostained for E-cadherin (**Fig. 2E**). There was a clear negative correlation between E-cadherin intensity at cell-cell junctions and the average RFP intensity of the two adjacent cells (**Figs. 2E-F, S2F**). Importantly, E-cadherin intensity at junctions between surface-derived and temporarily interior-located high-RFP cells (based on live-cell tracking) and their neighbors were indistinguishable from randomly sampled junctions between high-RFP cells (mostly at the surface) and their neighbors (**Figs. 2G, S2G**). Thus, we conclude that surface-derived cells maintain low E-cadherin expression when they navigate in the gland interior, which probably underlies their robust return to the surface (**Fig. 2B**).

### Accelerated branching of salivary glands upon basement membrane recovery from enzymatic disruption

Live imaging analysis and the E-cadherin expression patterns have led us to propose a model of salivary gland clefting based on the interplay between the basement membrane matrix and two cell types with distinct cell adhesion properties (**Fig. 2H**). In this testable model, provisional surface cells are first generated by proliferation of surface cells and temporarily stored in the interior domain to build up the “branching potential” or the relative abundance of interior-located low E-cadherin cells. These cells are then returned to the surface layer by cell sorting driven by differential cell-cell adhesion, reinsert between surface cells adhering weakly to each other, and use strong cell-matrix adhesions to remain adherent to the basement membrane. The expanded extra surface then drives folding of the surface epithelial sheet, causing clefting and new bud formation (**Fig. 2H**).

This model has an interesting prediction: if the number of stored interior proliferating cells could be increased, it might be possible to accumulate “branching potential” separate from actual branching. We tested this prediction by treating salivary glands with collagenase to disrupt the major basement membrane component collagen IV. While low concentrations of collagenase and antibody blocking of integrins inhibited salivary gland branching (**Fig. S3G-H**), high-concentration collagenase treatment caused existing epithelial buds to fuse together to revert branching (**Fig. 3A**) (Grobstein and Cohen, 1965; Rebustini et al., 2007). High-concentration collagenase virtually ablated collagen IV and greatly reduced laminin in basement membranes (**Fig. 3D**) without apparent effect on the surrounding mesenchyme (**Fig. 3E**). Importantly, we discovered greatly accelerated catch-up branching after collagenase washout (**Fig. 3A-C**), likely resulting from attaching of accumulated surface-originated post-proliferation cells to the restored basement membrane. We conclude that basement membrane disruption can uncouple surface expansion from the buildup of an interior pool of surface cell daughters. Basement membrane restoration enables rapid surface expansion and branching due to basement membrane anchorage and expansion of the surface epithelial sheet from this built-up interior pool of daughter cells.

**Figure 3.**
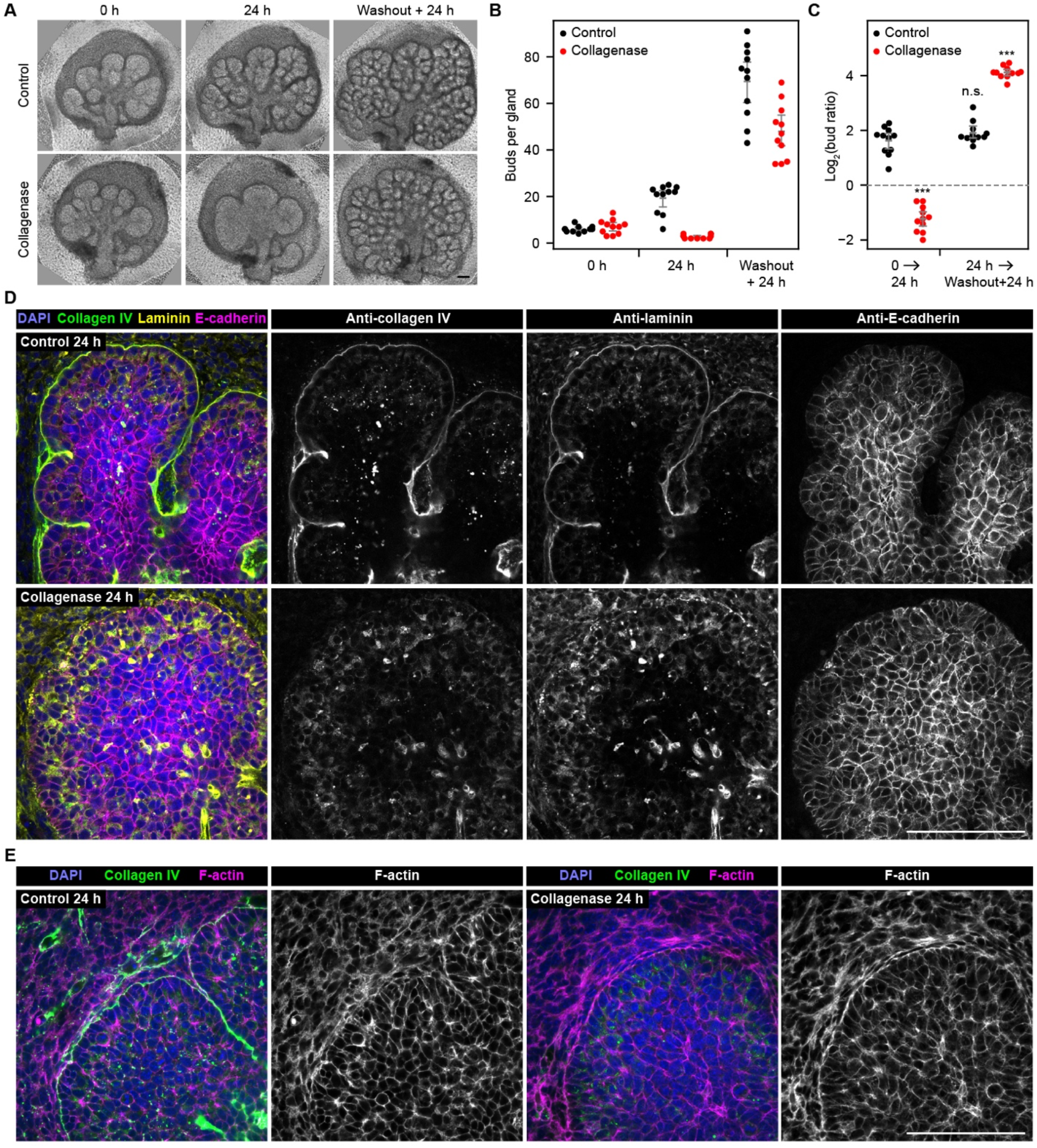
Accelerated branching of salivary glands upon basement membrane recovery from enzymatic disruption. (**A**) Phase contrast images of E12 + 1.5-day cultured salivary glands (0 h) treated for 24 hours with solvent control or 20 μg/mL collagenase (24 h), followed by another 24 hours after washout (washout + 24 h). (**B**-**C**) Plot of bud number per gland (B) or log2 bud ratio (C) over time. n=11 for each group. Error bars, 95% confidence intervals. ***, Tukey test p<0.001. n.s., not significant. All comparisons were to the first 24 hours of Control. (**D-E**) Confocal immunofluorescence images of control or collagenase treated glands at 24 h. Scale bars, 100 μm.

We next asked whether extra surface would be generated by comparing the shapes, sizes and cell proliferation rates of interior and surface-layer domains. We first derived numerical constraints for cell proliferation ratios (α) and geometric ratios (β) between interior and surface-layer domains of an inflating sphere (or oblate sphere) (**Fig. S3A-B**). An important assumption was compartmentalization between interior and surface domains, based on the observed clear separation of peripheral vs. interior epithelial cells (**Videos S3-4**) with robust surface return of surface-derived proliferated interior cells (**Fig. 2B**). This model predicts that extra surface will fold to maintain tissue integrity when the surface expands faster than the interior (Scenario II; **Fig. S3C**), as observed during salivary gland branching. We then estimated actual proliferation ratios (α) and geometric ratios (β) by immunostaining cell proliferation and morphology markers in branching salivary glands (**Fig. S3D-E**). We found that all data mapped to the parameter space permissive for surface folding (Scenario II; **Fig. S3F**), supporting our proposed model.

### Single-cell transcriptome profiling reveals spatial transcriptional patterns of the branching salivary gland epithelium

To explore regulatory mechanisms underlying differential cell adhesion properties among epithelial cells, we profiled single-cell transcriptomes of the E13 salivary gland epithelium by single-cell RNA sequencing (scRNA-seq). The 6,943 single-cell transcriptomes formed 7 main clusters with distinct marker genes (**Fig. 4A, D, E**). The cluster identities were assigned based on combinatory expression profiles of known marker genes, including the bud marker Sox10, duct marker Sox2, basal epithelial (outer bud and duct) marker Krt14 and the luminal (or inner) duct marker Krt19 (Lombaert and Hoffman, 2010; Szymaniak et al., 2017). We validated the bud enrichment of Sox10 expression and the duct enrichment of Sox2 expression by single-molecule RNA fluorescence in situ hybridization (smFISH) (**Fig. S4B-C**) (Raj et al., 2008; Wang, 2018). In addition, we identified Cldn10 as a marker with strong inner bud enrichment and found that its protein product Claudin 10 was indeed highly expressed in the inner bud (**Figs. 4D-E, S4E**). Although it might initially appear counter-intuitive, calculations based on gland dimensions indicate that there should be significantly more outer bud cells than inner bud cells (**Fig. S4A**).

**Figure 4.**
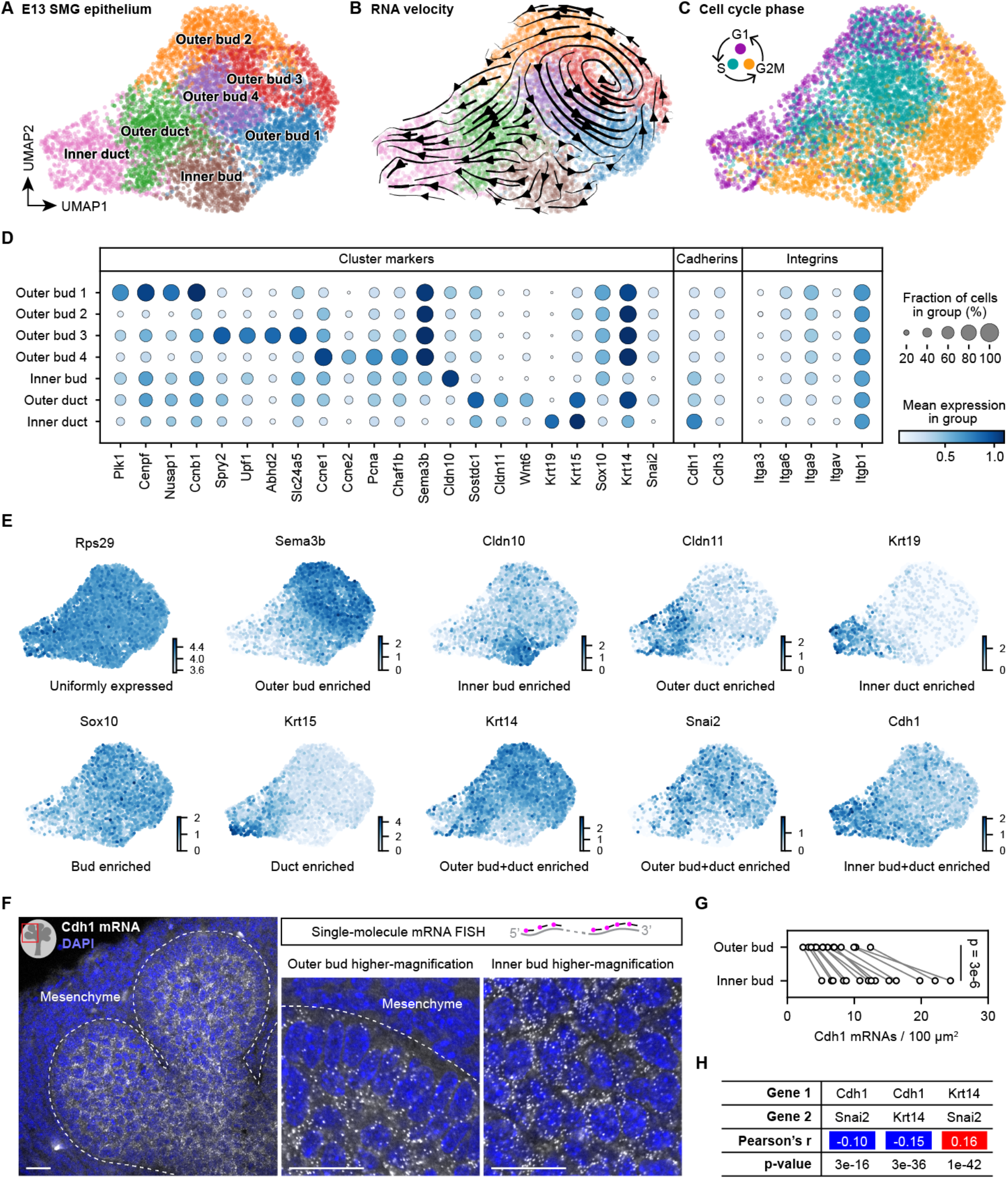
Single-cell transcriptome profiling reveals spatial transcriptional patterns of the branching salivary gland epithelium. (**A-C**) Scatter plots of 6,943 single-cell transcriptomes from the E13 mouse salivary gland epithelium shown in UMAP embedding and color coded by clusters (A and B) or cell cycle phase (C). Each dot represents one cell. Arrows in (B) indicate local RNA velocity estimated from unspliced and spliced transcripts of nearby cells. (**D**) Dot plot of selected cluster marker genes and cadherin and integrin genes. (**E**) Scatter plots of single-cell transcriptomes of E13 salivary gland epithelium in UMAP embedding and color coded by the expression level of indicated genes. (**F**) Confocal fluorescence images of Cdh1 mRNAs detected by single-molecule mRNA FISH in an E13 salivary gland. Each white dot is one Cdh1 mRNA molecule. (**G**) Plot of the Cdh1 mRNA density in outer or inner epithelial bud of E13 salivary glands. Measurements from the same image were connected by a line. p-value, paired two-sided t-test. (**H**) Table of the Pearson’s correlation coefficients and p-values between indicated genes. Blue and red shadings indicate negative and positive correlations, respectively. Scale bars, 20 μm.

To evaluate the dynamics of single-cell transcriptomes, we calculated RNA velocity (Bergen et al., 2020; La Manno et al., 2018), which predicts the future state of individual cells based on their unspliced and spliced mRNAs (**Fig. 4B**). This analysis revealed two prominent patterns: a cycling vector field covering all 4 outer bud clusters and a directional flow from the bud to duct clusters. The cycling vector field across outer bud clusters suggested outer bud cells were cell-cycling progenitors (Bergen et al., 2020), which was confirmed by the cell cycle phases of these cells (**Fig. 4C**). In fact, many marker genes of outer bud clusters were related to cell division or cell cycle regulation (**Fig. 4D**). On the other hand, the directional bud-to-duct flow suggested that some bud cells would differentiate into duct cells (**Fig. 4B**).

We next compared expression patterns of major cell adhesion genes. For cell-matrix adhesion, all prominently expressed integrin genes had comparable expression levels between outer bud and inner bud cells (**Figs. 4D, S4D**), indicating that integrin expression was not tightly regulated at the transcriptional level between these cells. For cell-cell adhesion, the E-cadherin gene Cdh1 showed clear enrichment in the inner bud and duct clusters compared to outer bud clusters (**Fig. 4D-E**), reminiscent of the expression pattern of E-cadherin protein (**Figs. 2C-E, S2F**). We confirmed this pattern of Cdh1 mRNA expression by smFISH (**Fig. 4F-G**). This alteration was specific, since expression of the P-cadherin gene Cdh3 showed no obvious enrichment in any cluster (**Fig. 4D**). Among the transcriptional factors involved in epithelial– mesenchymal transition (Stemmler et al., 2019), only Snai2 was prominently expressed in this tissue, and its expression pattern was negatively correlated to Cdh1 and positively correlated to Krt14 (**Fig. 4D, E, H**), suggesting a regulatory role for Snai2 in shaping the expression pattern of Cdh1, consistent with prior in vitro findings (Bolós et al., 2003).

### Reconstitution of epithelial branching morphogenesis using primary salivary gland epithelial cells

Next, we evaluated whether stratified epithelial branching could be reconstituted using primary salivary gland epithelial cells. Our lab had previously demonstrated self-assembly of dissociated salivary gland epithelial cells and partial primitive branching of self-assembled epithelial aggregates when dissociated cells were embedded in solidified high-concentration Matrigel (basement membrane matrix extract) (Kleinman et al., 1986) and cultured on a polycarbonate filter (Wei et al., 2007). The previous partially reconstituted branching might have been limited by epithelial cell attachment to the filter, and the reconstitution might be improved using low-attachment culture vessels. We tested several culture conditions and achieved optimal results using a 96-well ultra-low attachment plate, greatly reduced Matrigel supplement, and an enhanced recipe for organ culture media (Nakao et al., 2017). Under optimal conditions, we were able to recapitulate prominent branching morphogenesis of either isolated single epithelial buds or completely dissociated single epithelial cells with rates comparable to intact salivary gland culture (**Fig. 5**). Reconstituted branching from dissociated primary epithelial cells formed both end bud and duct structures **(Fig. 5E**). Thus, we conclude that key aspects of stratified epithelial branching morphogenesis can be reconstituted from primary epithelial cells without the mesenchyme. These findings suggested the possibility of attempting partial reconstitution of budding or branching morphogenesis using non-embryonic engineered cells.

**Figure 5.**
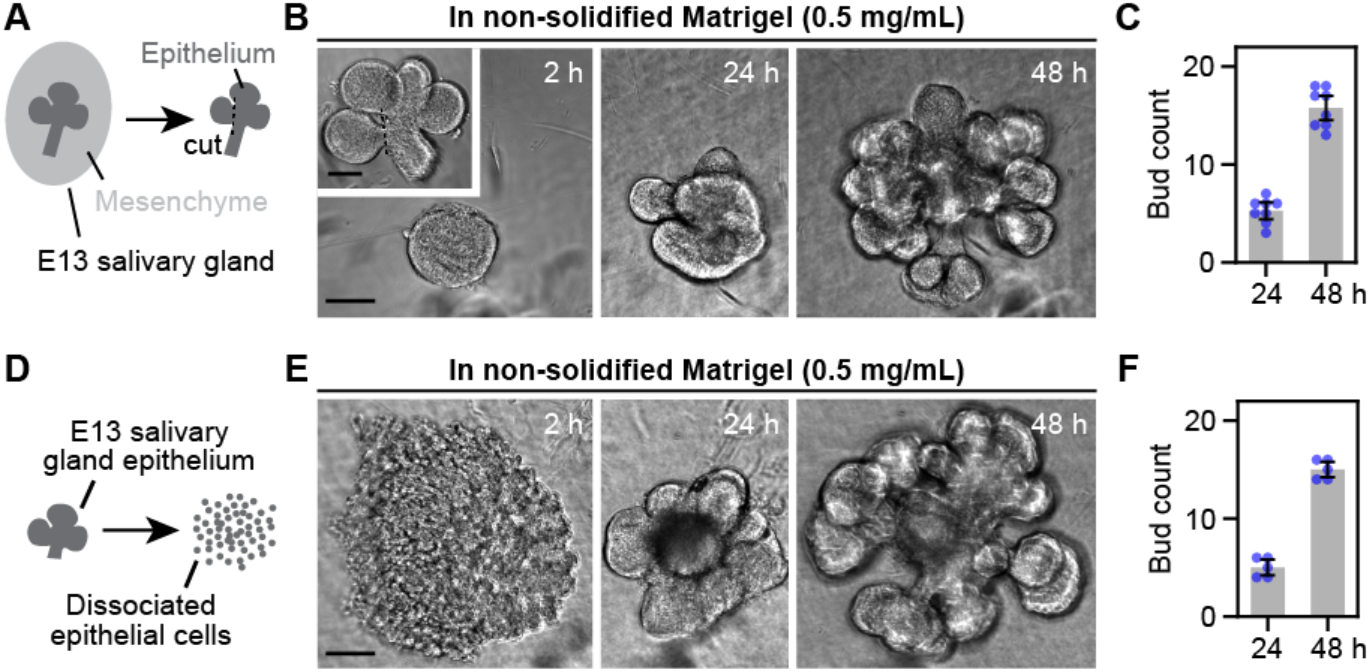
Reconstitution of epithelial branching morphogenesis using primary salivary gland epithelial cells. (**A, D**) Schematics of single-bud (A) and single-cell (D) isolation from E13 salivary glands. (**B, E**) Phase contrast images at indicated time points from single-bud (B) or single-cell (E) cultures. (**C, F**) Plots of bud number of single-bud (C) or single-cell (F) cultures. Error bars, 95% confidence intervals. Scale bars, 100 μm.

### Reconstitution of epithelial budding morphogenesis by engineering cell adhesion

Our model (**Fig. 2H**) suggests that the key initial process of stratified epithelial budding is driven by a population of cells with weak cell-cell adhesions plus strong cell-matrix adhesions.Reducing cell-cell adhesion can positively regulate branching morphogenesis in both mammary gland and embryonic pancreas (Nguyen-Ngoc et al., 2012; Shih et al., 2016), but whether it alone is sufficient to drive branching remained unknown. To address this question, we attempted to reconstitute epithelial branching by engineering cell adhesion. We chose the human adult colorectal adenocarcinoma cell line DLD-1 as a starting point, because DLD-1 expresses abundant E-cadherin and forms near-spherical spheroids without buds under low-attachment 3D culture conditions (Riedl et al., 2017). For sophisticated modulation of different cell adhesion molecules, we established a clonal DLD-1 cell line after introducing transgenes to enable CRISPR/dCas9-based inducible transcriptional repression and activation (Gao et al., 2016) (**Fig. S5A-B**). To stably express sgRNAs and simultaneously monitor their expression, we constructed lentiviral vectors co-expressing sgRNAs with bright nuclear fluorescent reporters (**Fig. S5C**).

To reduce cell-cell adhesion strength, we identified two Cdh1 sgRNAs that efficiently reduced E-cadherin expression levels after cells were treated with abscisic acid (ABA), a dimerizer used to recruit the KRAB transcriptional repression domain (**Figs. 6A, S5D-F**). Without ABA, sg1-Cdh1 had minimal effects, whereas sg2-Cdh1 reduced E-cadherin to ∼20% of controls (**Fig. S5D-F**), likely due to direct transcriptional blockade (Qi et al., 2013). The level of E-cadherin reduction could be titrated by ABA concentrations, approaching maximum reduction at 3 days for sg1-Cdh1 and 2 days for sg2-Chd1 (**Fig. S5G-H**). Inhibiting E-cadherin in DLD-1 resulted in only a moderate reduction of total β-catenin (**Fig. S5I**) and did not reduce cell proliferation or survival as reported for breast cancer cells (Padmanaban et al., 2019). Consistent with this, β-catenin in the cytoplasm and nucleus remain unchanged despite a severe loss from cell junctions upon E-cadherin downregulation (**Fig. S5J**). Importantly, we observed sorting out of low-E-cadherin cells in spheroid cultures of mixed sg-Control and sg-Cdh1 cells, suggesting E-cadherin reduction successfully lowered cell-cell adhesion strength (**Fig. S6A**).

**Figure 6.**
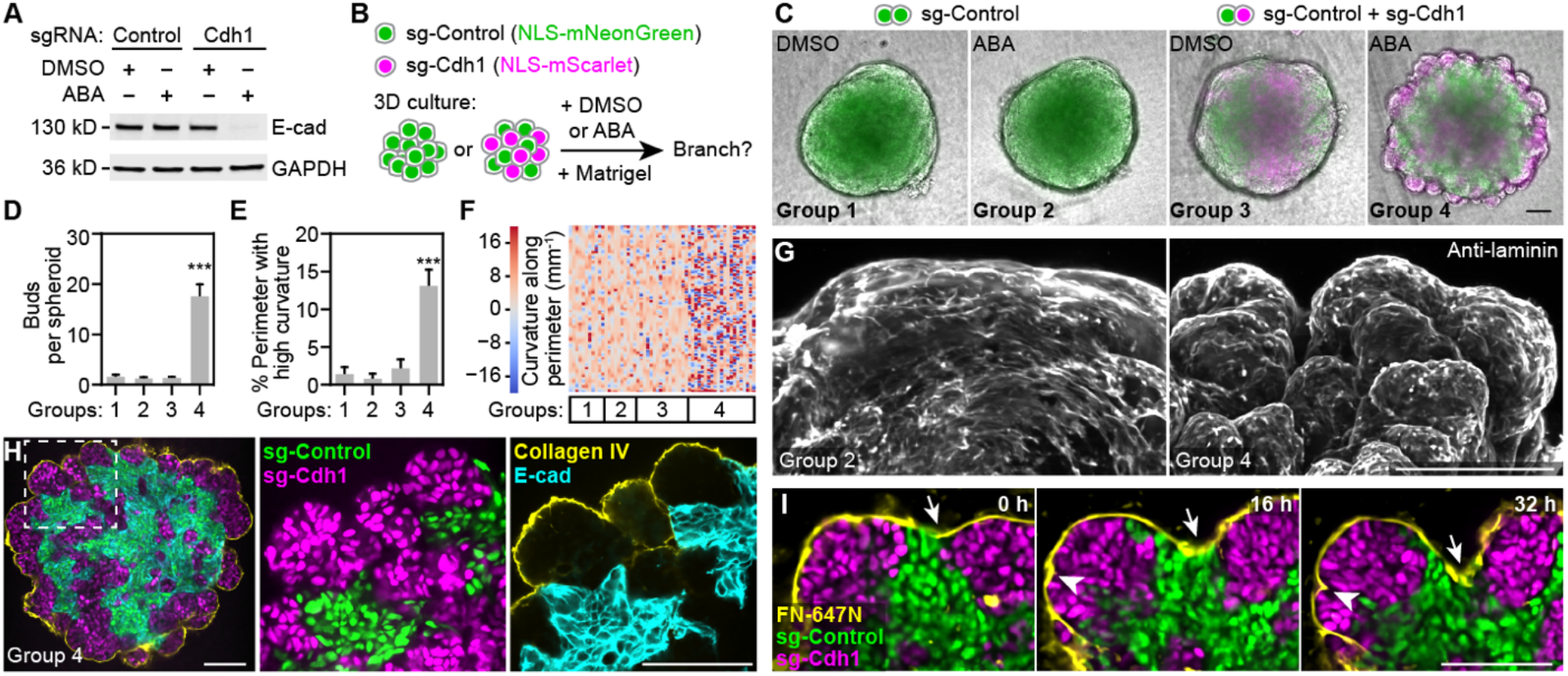
Reconstitution of epithelial budding morphogenesis using engineered cells. (**A**) Western blot of clonal Dia-C6 cells expressing Control (lacZ) or Cdh1 sgRNA treated with abscisic acid (ABA) or DMSO (vehicle). ABA is a dimerizer used to induce robust transcriptional repression in engineered cells. (**B**) Schematic of 3D spheroid cultures. (**C**) Merged phase contrast and epifluorescence images of spheroids from indicated experimental groups. (**D**-**E**) Bar plots of bud number or percentage of high-curvature perimeter length (|curvature|>20 mm^-1^). Error bars, 95% confidence intervals. ***, Tukey test p < 0.001. (**F**) Heatmap showing color-coded curvature along spheroid perimeters. Each column is one spheroid. Sample numbers in (C to F): n=11, 10, 16, 43 for groups 1-4 combining 2 independent experiments with similar results; only 21 randomly selected Group 4 samples were plotted in (F) to save space. (**G**) Maximum intensity projection of two-photon microscopy images of spheroids immunostained with laminin, a basement membrane marker. (**H**) Confocal images of a spheroid at the central slice. (**I**) Time-lapse confocal images of a branching spheroid. Atto-647N-labeled fibronectin was used to mark the basement membrane (yellow); arrows and arrowheads indicate clefts. Scale bars, 100 μm.

DLD-1 spheroids failed to spontaneously form a basement membrane (data not shown), a structure critical for salivary gland branching (**Fig. 3**). To induce basement membrane formation, we supplemented culture media with a non-solidified, low-concentration suspension of the basement membrane extract Matrigel. Strikingly, this led to robust budding morphogenesis in spheroid cultures containing sg1-Cdh1 or sg2-Cdh1 cells after ABA-induced E-cadherin reduction (**Figs. 6B-F, S6B-F; Video S6**). Importantly, a condensed layer of basement membrane had formed around the spheroids with high levels of the basement membrane components laminin and collagen IV (**Fig. 6G-H**). In spheroids containing both sg-Cdh1 and sg-Control cells, cells contacting the basement membrane were primarily sg-Cdh1 cells lacking E-cadherin expression (**Figs. 6H, S6H**). Furthermore, live-spheroid imaging revealed preferential outward expansion of contact surfaces between sg-Cdh1 cells (magenta) and the basement membrane (yellow), whereas contact surfaces between sg-Control cells (green) and the basement membrane was largely found at the cleft bottom (**Fig. 6H-I; Video S9**). Bud formation could also occur in spheroids containing only low-E-cadherin cells, but these spheroids were often flatter, and their buds were more amorphous (**Fig. S6G**), suggesting high-E-cadherin cells may play a structural role by forming a more robust spheroid core. To summarize the process of reconstituted budding morphogenesis, cells with experimentally reduced E-cadherin expression sorted out to the surface by differential cell-cell adhesion and then interacted with the basement membrane to promote budding as strong cell-matrix adhesions displaced weak cell-cell adhesions (**Fig. S6I**).

### Reconstituted epithelial budding depends on integrin-mediated cell-matrix adhesion

Our model predicts that strong cell-matrix interactions are required for reconstituted budding morphogenesis. We tested this by inhibiting cell-matrix interactions using a function-blocking β1-integrin antibody, which inhibited bud formation (**Fig. S7A**). In addition, enzymatic disruption of the basement membrane reverted budding, which could recover upon basement membrane reformation (**Fig. S7B**). We tested whether β1-integrin-dependent cell-matrix interactions were specifically required in low-E-cadherin cells using an Itgb1 sgRNA that efficiently reduced β1-integrin expression after ABA-enhanced transcriptional repression (**Fig. S7C-E**). Bud formation was completely blocked when sg-Itgb1 was expressed in low-E-cadherin cells or in all cells, but not when in only the high-E-cadherin cells (**Figs. 7A-C, S7F**), demonstrating that β1-integrin-dependent cell-matrix interactions were specifically required in low-E-cadherin cells for budding. Importantly, reducing E-cadherin or β1-integrin expression selectively inhibited cell attachment to E-cadherin extracellular domain or Matrigel-coated surfaces, respectively (**Fig. S7G-J**).

**Figure 7.**
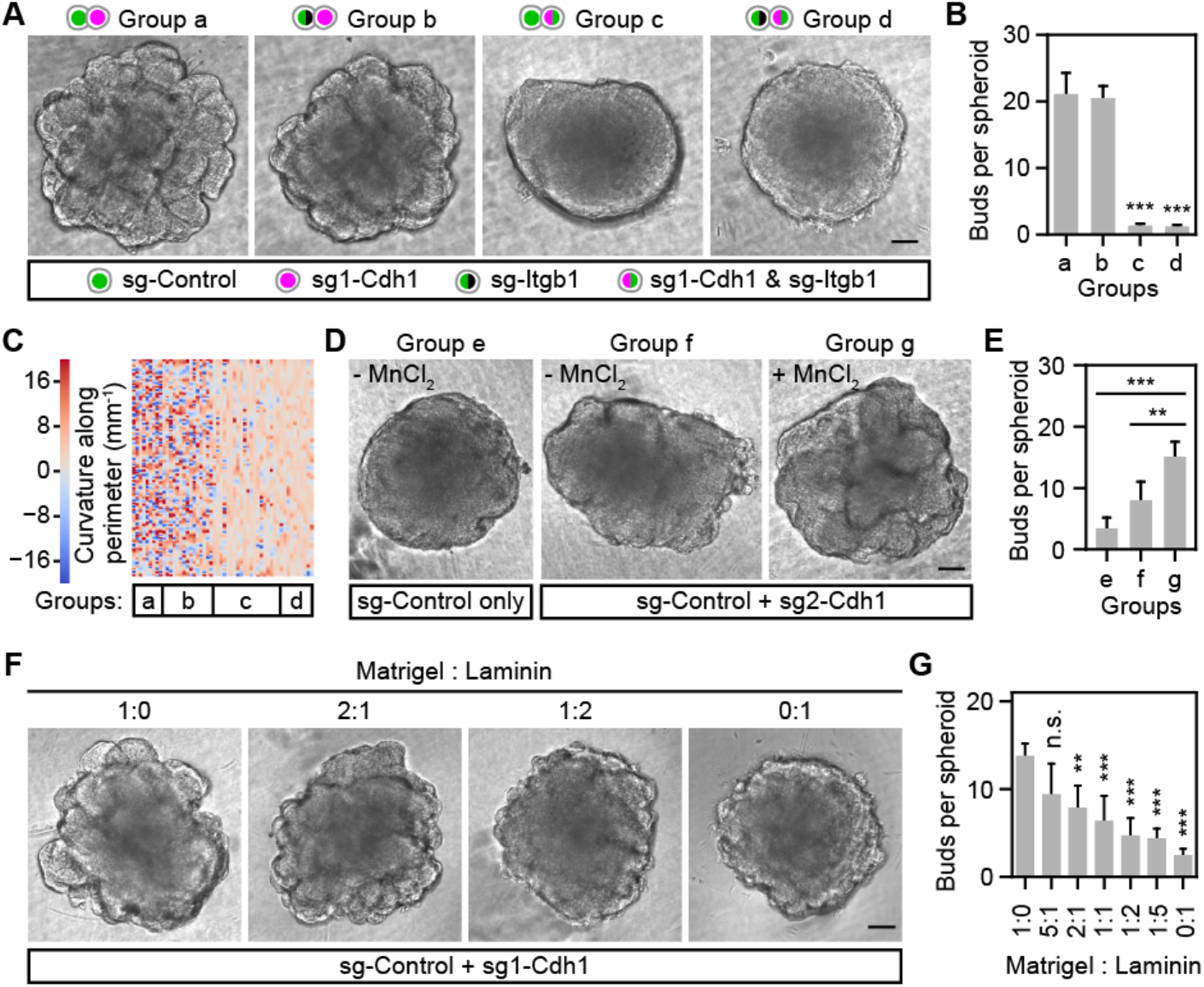
Reconstituted epithelial budding depends on integrin-mediated cell-matrix adhesion. (**A, D, F**) Phase contrast images of spheroids from indicated experimental groups. 50 μM MnCl2 was used to enhance integrin-mediated cell-matrix adhesion (D). ABA was added in all cultures to induce E-cadherin repression. Matrigel (A, D) or indicated ratios of Matrigel and laminin (F) were supplemented. (**B, E, G**) Bar plot of bud number per spheroid. (**C**) Heatmap showing color-coded curvature along spheroid perimeters. Each column is one spheroid. Sample numbers: n=9, 15, 20, 10, 10, 10, 20 for groups a-g from one of two independent experiments with similar results. n=10 for all groups in (G). Error bars in (B, E, G), 95% confidence intervals. **, ***, Tukey test p < 0.01 or p < 0.001. n.s., not significant. Scale bars, 100 μm.

We next tested whether enhancing cell-matrix adhesion strength could enhance budding. We capitalized on the low-level bud formation observed in mixed sg-Control/sg2-Cdh1 spheroid cultures without ABA, providing a sensitized assay (**Fig. S6C-F**). Using MnCl_2_ to enhance integrin-mediated cell-matrix adhesion strength (Bazzoni et al., 1995) produced a modest but definitive increase in bud formation (**Fig. 7D-E**). Thus, enhancing cell-matrix adhesion strength could enhance budding.

Finally, we tested whether changing the composition of supplemented matrix would affect the extent of reconstituted budding. Matrigel contains ∼60% laminin and ∼30% collagen IV (Kleinman et al., 1986). When mixing Matrigel with an increasing ratio of laminin to titrate down the ratio of collagen IV, we observed reduced budding morphogenesis (**Fig. 7F-G**), indicating the importance of an optimal matrix composition.

## DISCUSSION

Our work here has revealed the critical role of a specific combination of strong cell-matrix adhesions and weak cell-cell adhesions for driving budding morphogenesis of stratified epithelia, the key first step of branching morphogenesis. We have discovered that budding morphogenesis of stratified salivary gland epithelium is driven by the comparatively faster expansion and inward folding of a cryptic surface epithelial sheet, which we could visualize by single-cell tracking. Mechanistically, the expansion of this epithelial sheet is driven by the subsurface cell division and back-insertion of surface-derived daughter cells that retain weak cell-cell adhesions and re-establish strong cell-matrix adhesions to the basement membrane. Importantly, these two parameters were sufficient to successfully reconstitute budding morphogenesis of a stratified epithelium by experimentally reducing E-cadherin expression and inducing basement membrane formation to provide cell-matrix adhesion. Below, we discuss various roles of cell adhesion in epithelial morphogenesis and compare mechanisms used by single-layered vs. stratified epithelia in branching morphogenesis.

### Cell-matrix vs. cell-cell adhesion in epithelial morphogenesis

In addition to holding single cells together to form multicellular tissues, cell-matrix and cell-cell adhesions play important roles in epithelial morphogenesis. For example, differential cell-cell adhesion promotes cell sorting (Steinberg, 1963), whereas differing cell-matrix interactions between cell types can override the influence of cell-cell interactions to provide alternative self-organization strategies (Cerchiari et al., 2015). In both scenarios, tissues are driven toward states that minimize systemic interfacial energy by maximizing interfaces with stronger interactions. Following related biophysical principles, our work establishes that strong cell-matrix adhesions associated with weak cell-cell adhesions are sufficient to drive clefting and bud formation of a stratified epithelium.

Weak cell-cell adhesions of peripheral epithelial cells (and their descendants) are likely to play two important roles in promoting the expansion of the surface epithelial sheet. First, surface-derived epithelial cells that temporarily localize to the bud interior for cell division presumably rely on their inherited property of weak cell-cell adhesion to sort back out to the surface (**Fig. 2E-H**). Second, the weak cell-cell adhesions between the cells at the surface should allow returning daughter cells to intercalate between them to reach and engage with the basement membrane. Both processes would actualize the expansion of the surface epithelial sheet.

Conversely, the strong cell-matrix adhesions of peripheral epithelial cells to the basement membrane would both promote expansion of the surface epithelial sheet and maintain its integrity. In fact, when integrin-mediated cell-matrix adhesion is inhibited in salivary glands, surface cells frequently detach from the basement membrane and migrate into the bud interior, which inhibits branching (Hsu et al., 2013) (**Fig. S2H**). When the basement membrane is enzymatically disrupted in branched salivary glands, pre-existing epithelial buds fuse together (Grobstein and Cohen, 1965) (**Fig. 3A**), but cells expressing low E-cadherin still predominantly occupy the outer layers (**Fig. 3D**), presumably due to cell sorting based on differential cell-cell adhesion. In both normal branching and accelerated branching upon basement membrane recovery, the surface cell sheet expands because cells with low E-cadherin prefer to engage with the basement membrane rather than with other low E-cadherin cells (**Fig. 2H**). By increasing interfaces with stronger interactions, the epithelial cells and the basement membrane comprise a system proceeding towards a state of lower overall interfacial energy.

### Subsurface cell divisions in branching epithelia

Surface epithelial cells in the salivary gland epithelium predominantly divide in the subsurface layer after delaminating from the surface (**Fig. 2A**). Similar out-of-plane cell divisions have also been observed in the stratified embryonic pancreatic epithelium (Shih et al., 2016), as well as in single-layered embryonic lung and kidney epithelia (Packard et al., 2013; Schnatwinkel and Niswander, 2013). In single-layered epithelia, premitotic epithelial cells delaminate from the epithelial sheet, complete cell divisions in the lumen, and then reinsert into the epithelial sheet. In stratified epithelia, premitotic surface epithelial cells delaminate from the surface and complete cell divisions in the bud interior, which is packed with cells. Previously, the crowded cell environment made tracking of surface-derived daughter cells challenging in stratified epithelia, although some isolated examples of cell return to the surface were observed (Hsu et al., 2013; Shih et al., 2016). We establish here that all of the surface-derived daughter cells in the salivary gland epithelium unequivocally return to the surface, a conclusion made possible by our successful development of live-organ imaging strategies for high-resolution imaging and tracking of individual cells throughout the entire developing epithelium over an extended period of time.

It is not clear what drives the delamination of surface epithelial cells before they divide in this and other biological systems. We speculate it could be due to mitotic cell rounding and cell crowding in the surface layer. It is well-known that mitotic cells round up due to outward osmotic pressure and inward contraction of the actomyosin cortex (Stewart et al., 2011). When cells round up before mitosis, the surrounding cells in the crowded surface cell sheet will likely extrude them from the layer, since they all compete for attachment to the limited basement membrane surface area. Since the basement membrane provides a thin but dense barrier to cell migration, surface cells are pushed into the mass of jostling cells of the subsurface layer. An additional contributor might potentially be some direct pulling force from interior cells adhering to the surface cells, which is usually counterbalanced by the tight adherence of surface cells to the basement membrane. It might be interesting to test experimentally various possible mechanisms underlying the characteristic out-of-plane cell divisions of developing epithelial tissues.

### Reconstitution of budding using both primary and engineered cells

We have demonstrated reconstitution of epithelial morphogenesis using the two complementary systems of primary salivary gland epithelial cells and engineered DLD-1 cells. While 3D cultures from primary epithelial cells recapitulate both budding and ductal morphogenesis (**Fig. 5**), 3D cultures of engineered DLD-1 cells recapitulate just the first key steps of clefting and budding morphogenesis (**Figs. 6-7**).

A future direction could be to identify novel regulatory mechanisms of ductal morphogenesis to attempt reconstitution of full branching morphogenesis of a stratified epithelium using engineered cells. 3D cultures from primary epithelial cells can be used for perturbation studies to identify candidate regulatory molecules, while 3D cultures from engineered DLD-1 cells — a vastly different cell line — can enable testing of sufficiency, as shown in this study.

### Branching morphogenesis of stratified vs. single-layer epithelia

With the new insight here that the key first step of stratified epithelial branching can be conceptualized as folding of an expanding surface cell sheet, our findings reveal hidden similarities between the seemingly discrepant branching mechanisms used by single-layered and stratified epithelia. Both can now be conceptually understood as buckling of an epithelial sheet (Nelson, 2016), except that the surface cell sheet in a stratified epithelium is more cryptic until visualized by tracking cell dynamics.

Buckling of the two types of epithelial sheets is shaped by different constraints imposed by their surrounding tissues. Outside the basement membrane of both types of epithelial sheets, there is a mesenchyme consisting of both cells and extracellular matrix. However, while the epithelial sheet in a stratified epithelium is directly attached to an inner cell core, the epithelial sheet in a single-layered epithelium encloses a lumen filled with fluid. As a result, buckling of a single-layered epithelium is primarily constrained by the surrounding mesenchyme, as described in mouse embryonic lungs and intestines (Goodwin et al., 2019; Hughes et al., 2018; Kim et al., 2015). In contrast, buckling of the surface epithelial sheet in a stratified epithelium is constrained by both the surrounding mesenchyme and the interior epithelium. In the stratified salivary gland epithelium, the role of the interior epithelium seems to be more dominant than that of the surrounding mesenchyme. In our mesenchyme-free cultures of primary salivary gland epithelial cells in non-solidified, low-concentration Matrigel, budding morphologies are very similar to those of intact salivary gland cultures containing mesenchyme (compare buds in **Fig. 5** with **Figs. 1B, 3A**). In fact, as demonstrated in our simplified model, preferential expansion of a surface layer attached to an inner cell core alone is sufficient to drive the folding of the surface layer (**Fig. S3C**).

In conclusion, our study establishes the concept of the critical role of a specific combination of strong cell-matrix adhesion and weak overall cell-cell adhesion of peripheral epithelial cells for the expansion and buckling of a cryptic surface epithelial sheet, which in turn drives budding morphogenesis of a stratified epithelium. We anticipate that this unifying view of branching morphogenesis as buckling of an epithelial sheet will facilitate development of unifying physical models of branching morphogenesis that encompasses both single-layered and stratified epithelia.

## ACKNOWLEDGMENTS

We thank all members of the Yamada Laboratory for helpful discussions, J.W. Collins, D.A. Cruz Walma, A.D. Doyle, S.S. Nazari, D. Wu (NIDDK), J. Lu (Shandong University), A. Desai (UCSD), R.A. Green (UCSD), and K. Oegema (UCSD) for critical reading of the manuscript, A.D. Doyle from the NIDCR Imaging Core for assistance in microscopy, R. Sekiguchi for sharing tissue dissociation protocols, Z. Wei, D. Martin and R. Morell from the NIDCD/NIDCR Genomics and Computational Biology Core for scRNA-seq, L. Zhang, G.T. McGrady and E. Stregevsky from the NIDCR Combined Technical Research Core for Sanger sequencing and cell sorting, the NIDCR Veterinary Resource Core for mouse care, S. Qi (Stanford University), F. Zhang (MIT), and D. Trono (EPFL) for plasmids, M.P. Hoffman (NIDCR), E. Fuchs (Rockefeller University) and A.J. Ewald (JHU) for mice. Image analysis was performed in part using the NIH High Performance Computing system. This work was supported by the NIH Intramural Research Program (NIDCR, ZIA DE000525). S.W. was supported in part by an NIDCR K99 Pathway to Independence Award (K99 DE27982).

## AUTHOR CONTRIBUTIONS

S.W. and K.M.Y. conceptualized the project. S.W. designed experiments with useful input from K.M. and K.M.Y.. S.W. and K.M. performed all experiments. S.W. performed data analysis with useful input from K.M. and K.M.Y.. All authors contributed to data interpretation. S.W. and K.M.Y. wrote the manuscript with useful input from K.M.. K.M.Y. acquired funding and supervised the project.

## DATA AND CODE AVAILABILITY

Single-cell RNA sequencing data has been deposited in GEO (accession number GSE159780 for public access upon publication). Data for reproducing all other plots in this study are available in Figshare (reserved DOI for public access upon publication: 10.35092/yhjc.12145626). Raw data that support the findings of this study are available from the corresponding author upon reasonable request. Customized scripts and usage instructions are available from Github: https://github.com/snownontrace/public-scripts-Wang2020-branching-morphogenesis.

## LEAD CONTACT

Further information and requests for resources and reagents should be directed to and will be fulfilled by the Lead Contact, Kenneth M. Yamada.

**Figure S1 (Related to Figure 1).**
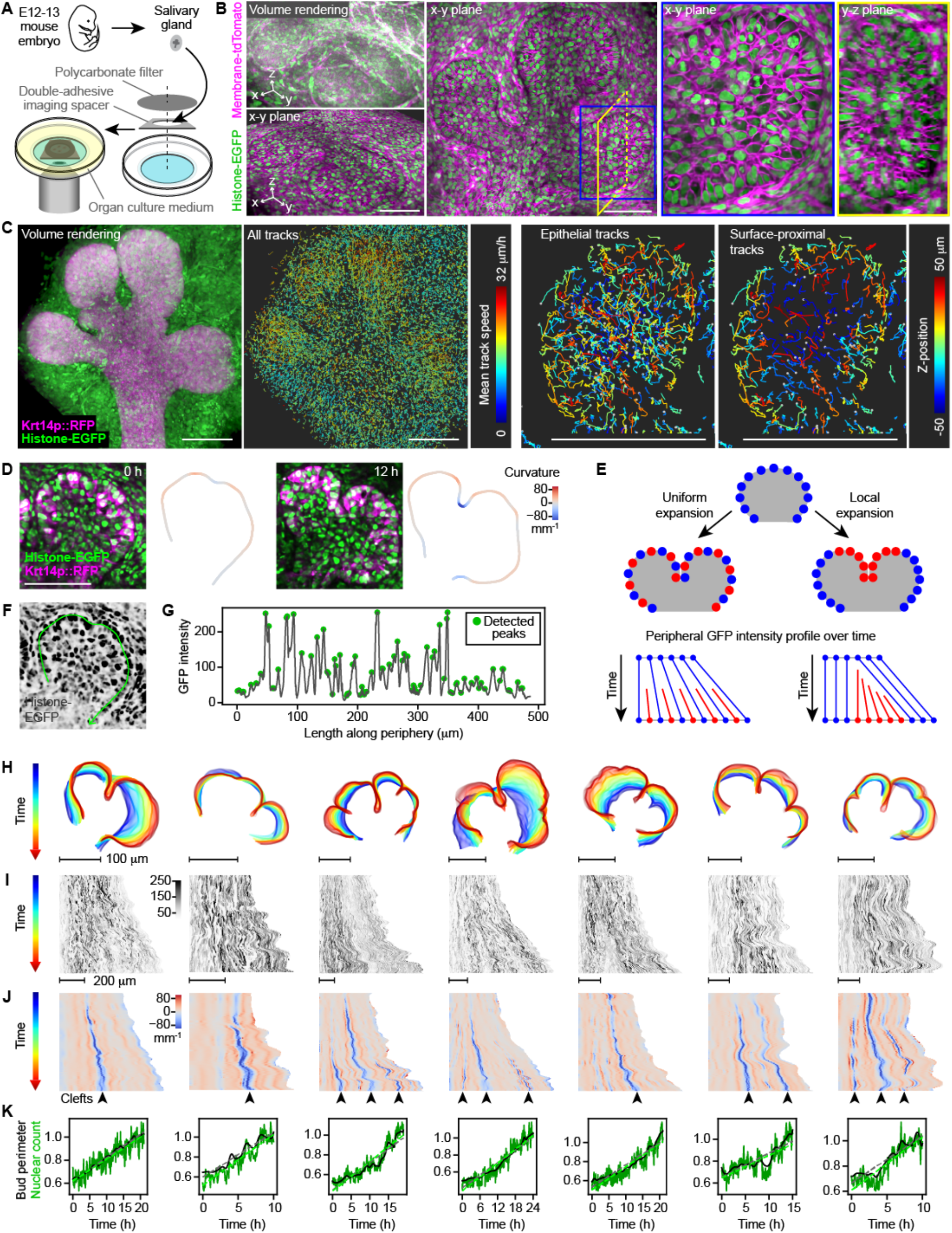
Live-imaging, cell tracking and surface expansion analysis of mouse embryonic salivary glands. (**A**) Schematic describing sample preparation for live imaging of wholemount mouse embryonic salivary glands. (**B**) Different views of a volumetric imaging data set at the first time point. (**C**) Views of confocal image and cell tracks of another volumetric imaging data set. (**D**) Two-photon microscopy images and epithelial surface outline color-coded by local curvature of an E13 transgenic mouse salivary gland. (**E**) Schematics of uniform vs. local expansion of the epithelial surface and their predicted peripheral GFP intensity profile over time. (**F**) Inverted histone-EGFP image from the same image in (D) at 0 h. The green curved line indicates the positions where peripheral GFP intensities were sampled. (**G**) Plot of GFP intensity along the green curve in (F). Green dots indicate automatically detected peaks used to quantify nuclear counts. (**H**-**K**) Outlines of the epithelial surface at the middle slice (H), heatmaps of the peripheral GFP intensity (I) and curvature (J), and plots of the bud perimeter (K; black) and peripheral nuclear count (K; green) over time. Dashed lines in (K) indicate fitted linear models. Scale bars, 100 μm.

**Figure S2 (Related to Figures 2 and 3).**
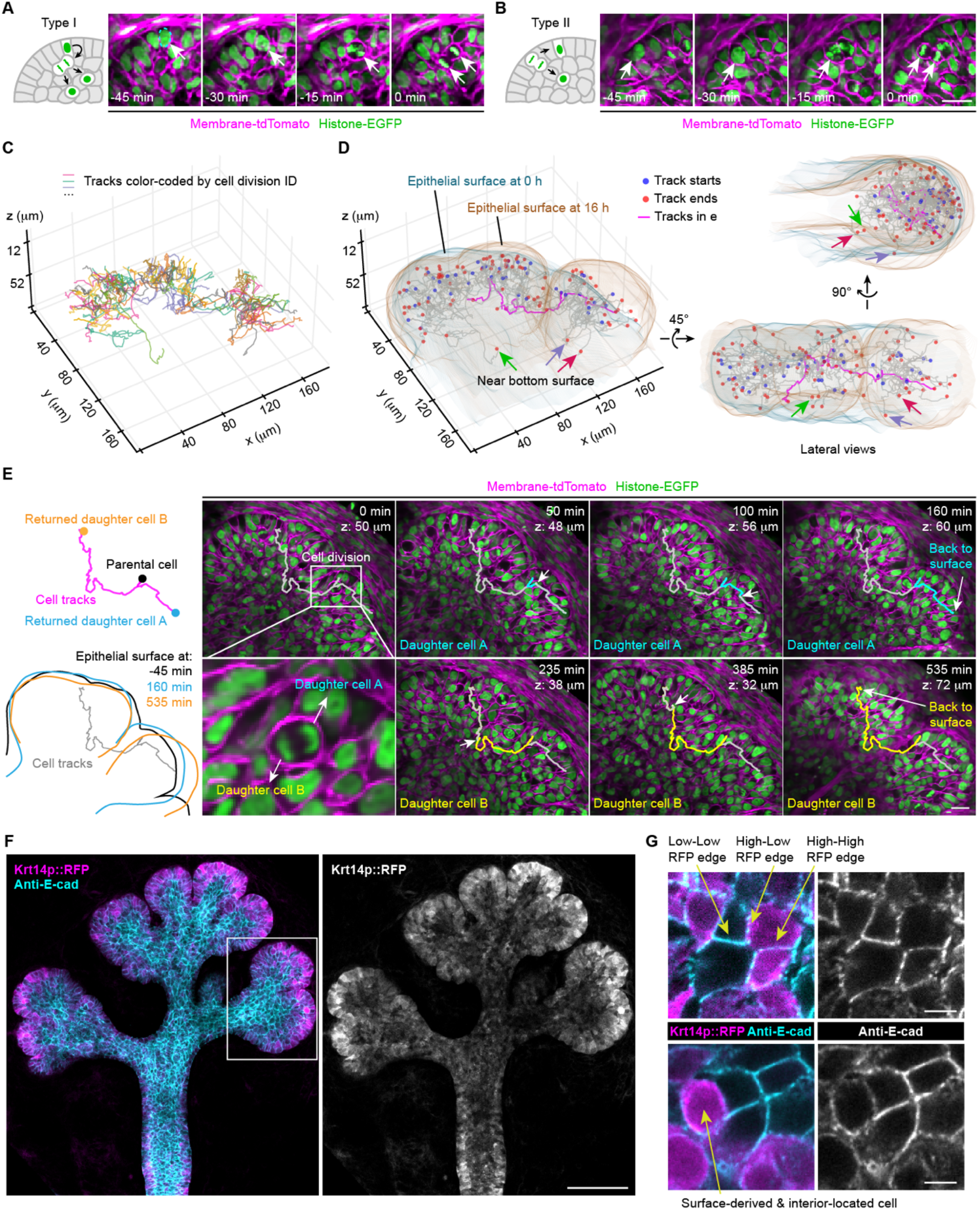
Robust surface return of interior daughter cells after cell division of surface epithelial cells. (**A**-**B**) Schematics and two-photon microscopy image sequences showing the two observed types of surface cell division. The 0 min images are the same as those in Fig. 2A. (**C**-**D**) 3D plots of cell tracks following 42 surface cell divisions and the surface return of their daughter cells. (**E**) Schematics (left) and snapshots of time-lapse two-photon images (right) following a cell division and tracks of both daughter cells highlighted in (D) and (E). White arrows point to tracked cells. (**F-G**) Confocal images of a transgenic Krt14p::RFP salivary gland immunostained for E-cadherin. White boxed region is the same as shown in Fig. 2E. Scale bars in (A, B, E), 20 μm; (F), 100 μm; (G), 5 μm.

**Figure S3 (Related to Figures 2 and 3).**
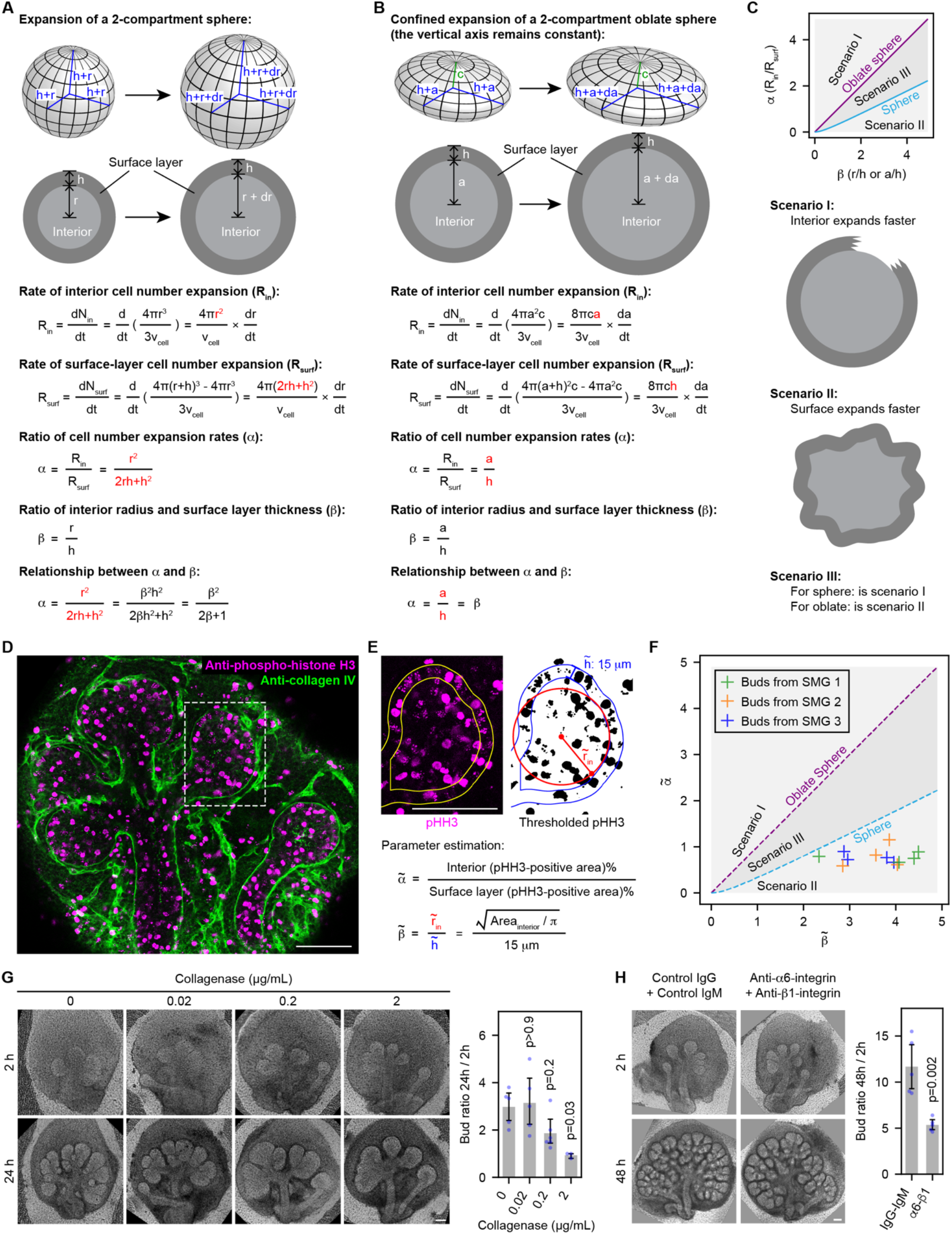
A simplified numerical model of stratified epithelial branching. (**A**-**B**) Schematics and mathematical derivation of the relationship between the parameters α and β during the expansion of a 2-compartment sphere (A) or an oblate sphere (B). Here α is the ratio of cell number expansion rates (interior vs. surface layer), whereas β is the ratio of interior radius vs. surface layer thickness. Important assumptions include: 1. Average cell volume remains constant. 2. Surface layer thickness remains constant. 3. The sphere or oblate sphere is filled with cells without spaces between cells. 4. The surface layer and interior compartments do not intermix. (**C**) Upper: plot of α vs. β. In the case of a sphere or an oblate sphere, α and β must fall on the turquoise or the purple line. Lower: illustrations of the three possible scenarios of parameter combination. (**D**) Confocal immunofluorescence image of an E13 salivary gland. Anti-phospho-histone H3 (pHH3) marks mitotic cells, and anti-collagen IV marks the basement membrane. (**E**) Upper: images illustrating the thresholding of the pHH3 signal. The outer boundary was automatically determined from anti-E-cadherin staining (not shown), which was shrunken by 15 μm to obtain the interior boundary. Lower: equations showing how estimated α and β were determined. (**F**) Plot of estimated α and β pairs for 12 epithelial buds from 3 submandibular salivary glands (SMG). Note that all parameter pairs fall within the Scenario II domain. (**G**) Phase contrast images (left) and bud ratio plot (right) of E13 salivary glands treated with indicated concentrations of collagenase. p-values, Tukey test compared to the 0 μg/mL group. (**H**) Phase contrast images (left) and bud ratio plot (right) of E13 salivary glands treated with control IgG and IgM or a combination of α6-integrin and β1-integrin antibodies. p-values, two tailed t-test. Scale bars, 100 μm.

**Figure S4 (Related to Figure 4).**
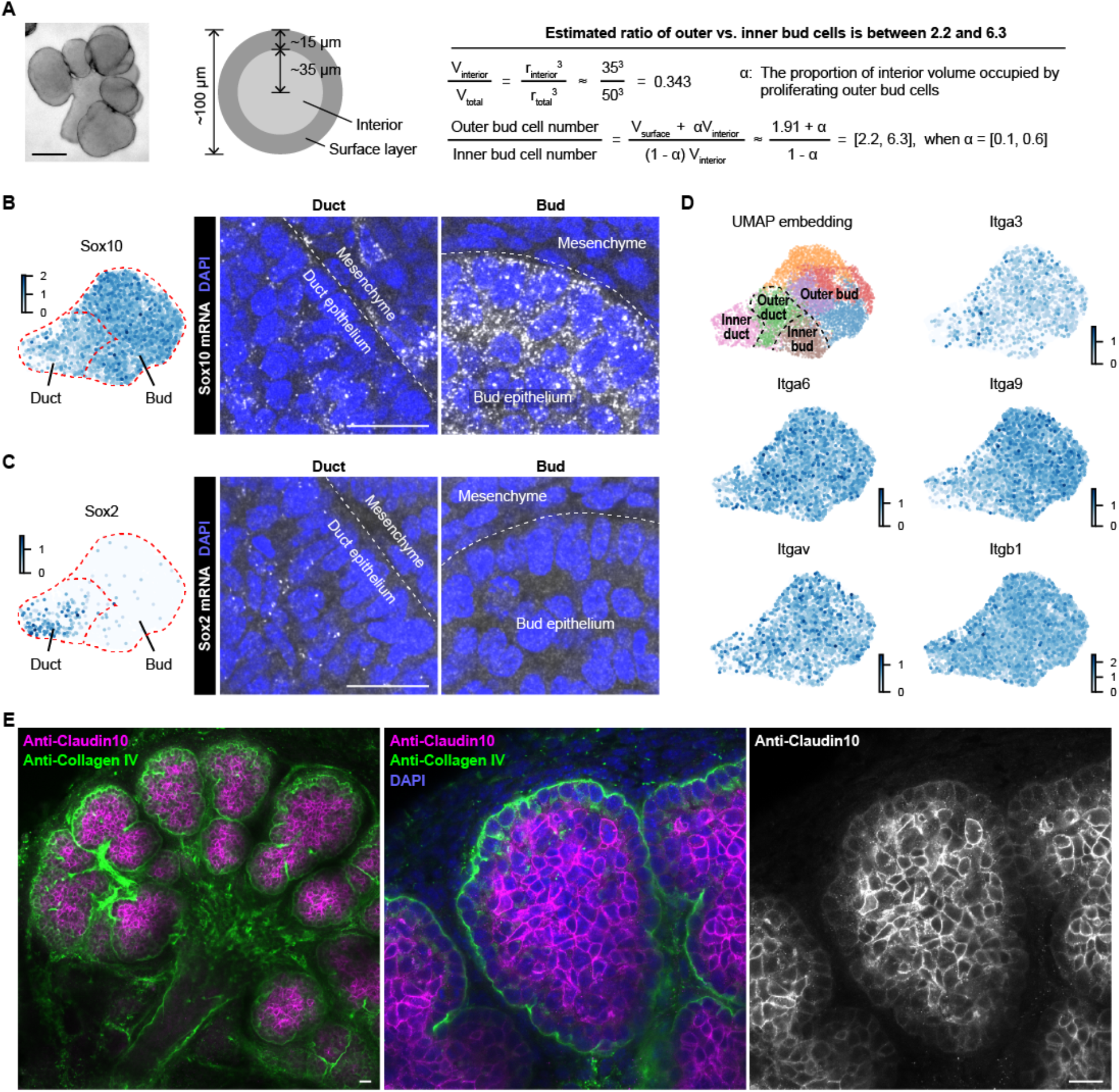
Expression patterns of selected marker genes from the scRNA-seq data set. (**A**) The estimated ratio of numbers of outer bud cells vs. inner bud cells in E13 salivary gland epithelium is between 2.2 and 6.3. (**B-C**) Scatter plots in UMAP embedding (left) and confocal images of smFISH staining (right) of Sox10 (B) or Sox2 (C). (**D**) Scatter plots in UMAP embedding color coded by clusters or indicated integrin genes. (**E**) Confocal immunofluorescence images highlighting an inner bud marker Claudin10 (encoded by the Cldn10 gene). Scale bar in (A), 100 μm; (B, C, E), 20 μm.

**Figure S5 (Related to Figure 6).**
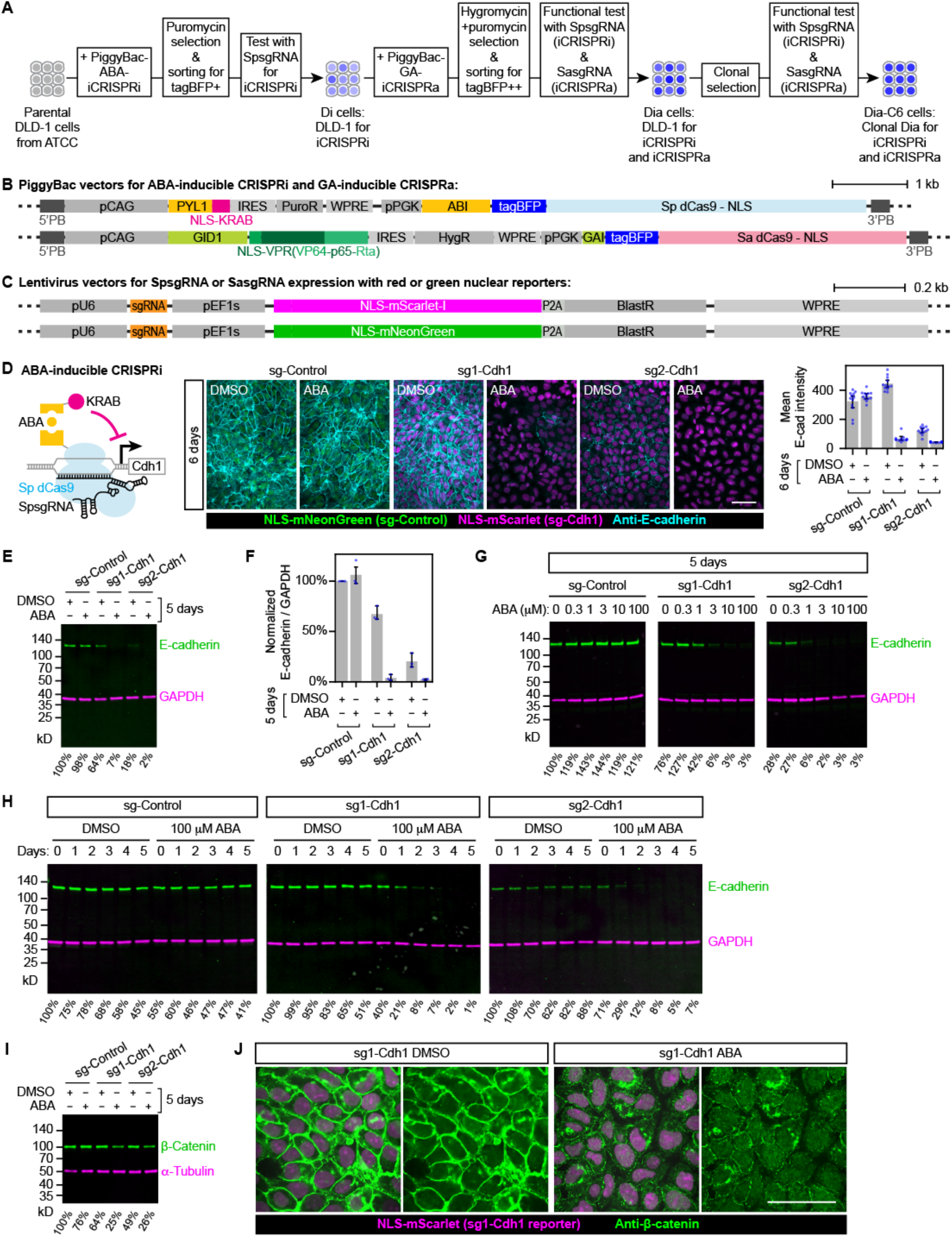
Establishing a clonal cell line for inducible modulation of cell adhesions. (**A**) Schematics illustrating engineering of a clonal Dia-C6 cell line from DLD-1. Dia cells stably express 4 transgenes that enable inducible transcriptional activation or repression. (**B**) Schematics of 2 PiggyBac vectors expressing 4 transgenes for inducible transcriptional modulation. (**C**) Schematics of lentivirus vectors used to deliver guide RNAs for transcriptional modulation. (**D**) Left: schematic of the ABA-inducible transcriptional repression system. Middle: maximum intensity projection of confocal images of cells expressing indicated sgRNAs. Green and magenta nuclei indicate cells expressing sgRNAs. Right: plot of mean E-cadherin intensities of indicated experimental conditions. n=12 fields of view for each group. (**E, G, H, I**) Western blots showing the expression level of E-cadherin or β-catenin under the indicated experimental conditions. Percentage levels under each lane are normalized ratios of E-cadherin to GAPDH or β-catenin to α-tubulin intensities of that lane. Normalizations are versus the control group (sg-Control or Day 0, DMSO) from the same blot. (**F**) Plot of normalized ratios of E-cadherin to GAPDH intensities of Dia cells expressing sg-Control, sg1-Cdh1 or sg2-Cdh1 treated with DMSO or ABA for 5 days. Normalizations are versus the control group (sg-Control, DMSO) for each of the 3 biological replicates. (**J**) Confocal images of β-catenin immunofluorescence. Error bars in (D, F) are 95% confidence intervals. Scale bars, 100 μm.

**Figure S6 (Related to Figure 6).**
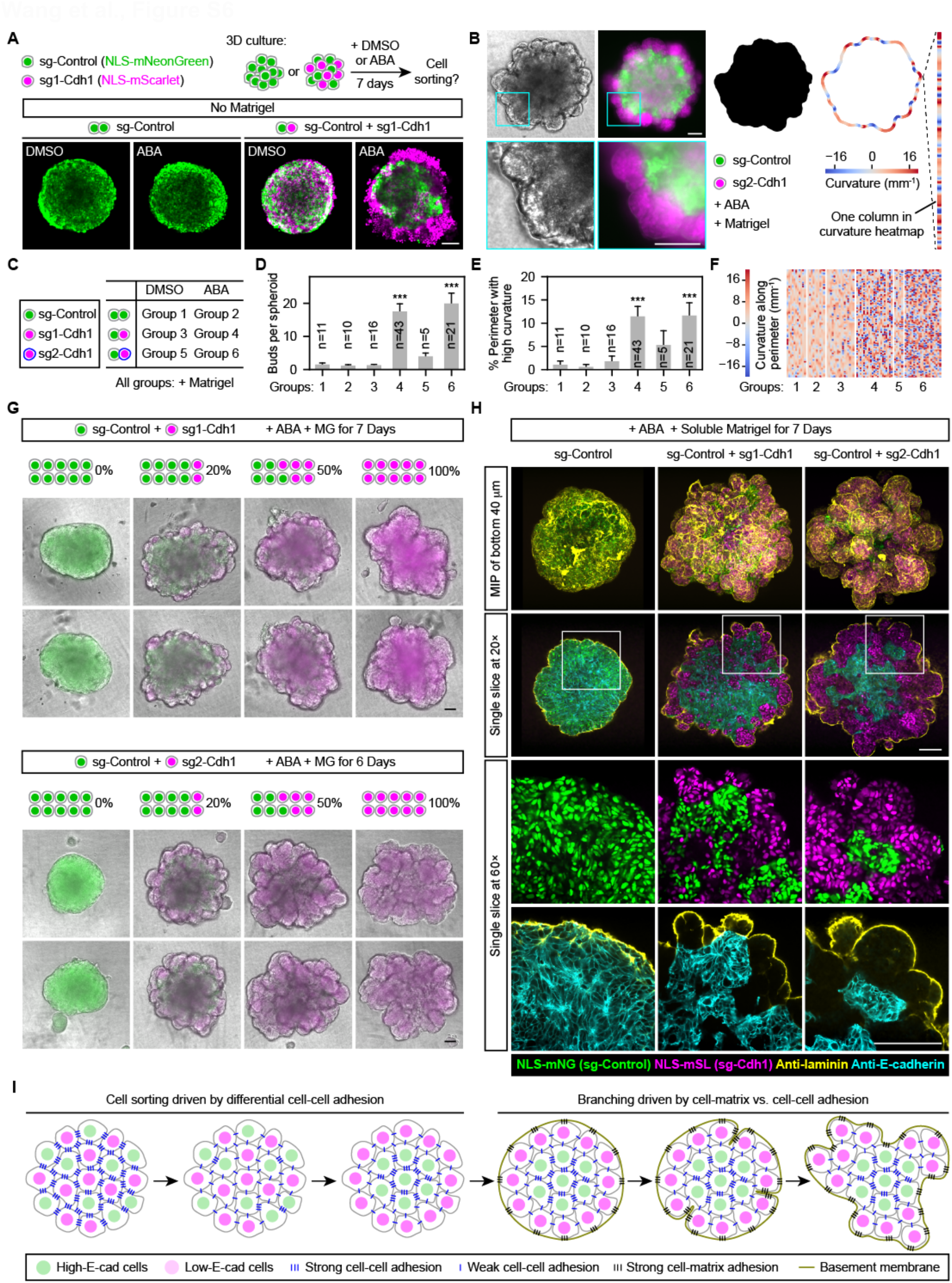
Reducing cell-cell adhesion and inducing basement membrane formation is sufficient to induce budding morphogenesis of a stratified epithelium. (**A**) Top: schematics of the experimental design. Below: confocal fluorescence images of spheroids for indicated experimental conditions. (**B**) Left: phase contrast and epifluorescence images of a spheroid. Right: binary mask image segmented from merged green and red fluorescence, and its boundary outline color-coded by the local curvature. Each column of curvature heatmaps represents a straightened outline color-coded by curvature. (**C**) Schematic and table showing experimental conditions of 6 groups. (**D**-**E**) Bar plots of bud number or percentage of high-curvature perimeter length (|curvature|>20 mm^-1^). ***, Tukey test p<0.001 compared to Group 1. (**F**) Heatmap showing color-coded curvature along spheroid perimeters. Each column is one spheroid. Only 21 of 43 Group 4 samples were included in the heatmap to save space. (**G**) Merged phase contrast and epifluorescence images of spheroids using two different sgRNAs under the indicated experimental conditions with different mixing ratios of control and E-cadherin downregulated cells. (**H**) Confocal images of spheroids from indicated experimental conditions immunostained with E-cadherin and laminin (yellow; a basement membrane marker). MIP, maximum intensity projection. (**I**) Schematics illustrating the cell sorting and branching of a stratified epithelium with mixed populations of high vs. low E-cadherin-expressing cells. Scale bars, 100 μm.

**Figure S7 (Related to Figure 7).**
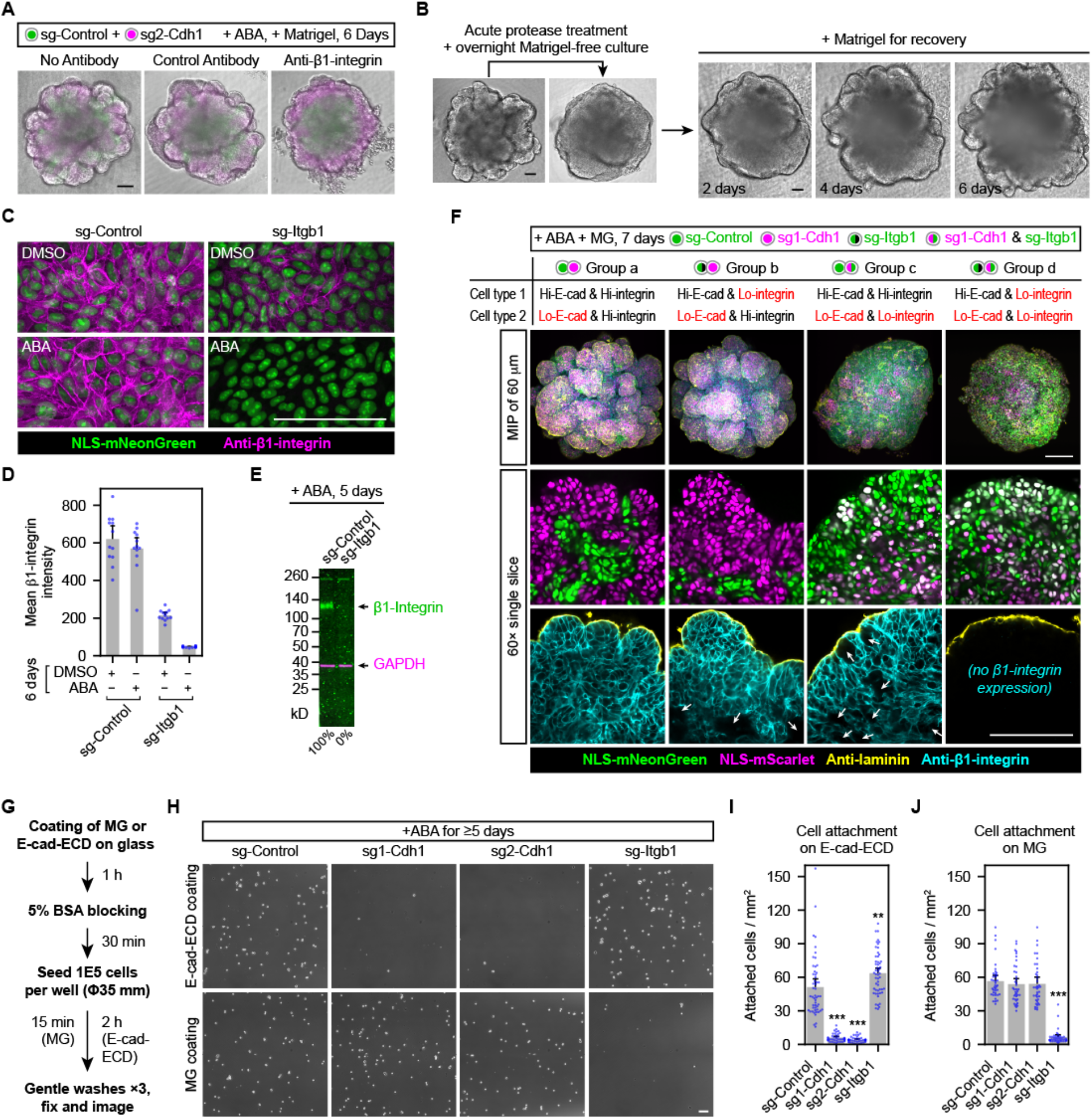
Reconstituted epithelial budding requires β1-integrin-mediated cell-matrix adhesion. (**A**) Merged phase contrast and epifluorescence images of spheroids from indicated experimental conditions. The α5-integrin antibody (clone mAb16) was used as a species-matched control antibody for the β1-integrin blocking antibody (clone mAb13). n=6 from 2 independent experiments. (**B**) Phase contrast images of budded spheroids (sg2-Cdh1 + sg-Control, ABA + Matrigel for 7 days) treated with dispase and recovered for 6 days after washout. n=5. (**C**) Maximum intensity projection of confocal immunofluorescence images of 2D-cultured cells expressing indicated sgRNAs. Green nuclei indicate cells expressing the sgRNAs. (**D**) Plot of mean β1-integrin intensities under the indicated experimental conditions. n=12 fields of view per group. (**E**) Western blot showing expression levels of β1-integrin under indicated experimental conditions. Percentage levels under each lane are normalized ratios of β1-integrin to GAPDH intensities of that lane. (**F**) Confocal images of spheroids under indicated experimental conditions. Note that green nuclei here indicate either sg-Control or sg-Itgb1 expression depending on the experimental group. White arrows point to cells that lost their β1-integrin expression. MIP, maximum intensity projection. (**G**) Schematics illustrating experimental procedures for the cell attachment assays. MG, Matrigel; E-cad-ECD, E-cadherin extracellular domain. (**H**) Phase contrast images of attached cells under indicated experimental conditions. (**I-J**) Plots of attached cells under indicated experimental conditions. n=52, 39, 39, 52 fields of view for groups in (I). n=39 fields of view per group in (J). Each experimental group had 3 or 4 biological replicates, and each had 13 fields of view. The full set of experiments was done twice with similar results. Scale bars, 100 μm.

## SUPPLEMENTAL VIDEO LEGENDS

**Video S1 (Related to Figure 1). Volumetric time-lapse video of a branching embryonic salivary gland**. Salivary gland from a 13-day transgenic mouse embryo expressing membrane-tdTomato and histone-EGFP. Time-lapse volumetric two-photon microscopy images were acquired at 2-μm z steps over a 100 μm z range at 5-minute intervals for 20 hours. The first half of the video shows different views of the imaging volume at the first time point. The second half of the video shows time-lapse sequences of the middle x-y plane (left) or 3D rendering of the middle 40 μm-thick volume of a branching epithelial bud (right).

**Video S2 (Related to Figure 1). Dynamics of surface-layer epithelial cells highlighted by KikGR photoconversion**. Salivary gland from a 13-day transgenic mouse embryo expressing the photoconvertible fluorescent protein KikGR. Native KikGR has green fluorescence, which converts to red fluorescence after blue light excitation. Magenta color indicates photoconverted KikGR.

**Video S3 (Related to Figure 1). Surface-proximal 3D cell migration tracks**. Videos show only epithelial cell tracks for which the closest distance to the gland surface was ≤ 15 μm. White dots indicate cell centroids, and the tracks shown at each time point represent cell migration trajectories spanning the previous 2 hours. Tracks are color coded by mean track speed (left) or the z position (right).

**Video S4 (Related to Figures 1 and 2). Different views of a branching epithelial bud from a developing salivary gland**. Salivary gland from a 13-day transgenic mouse embryo expressing histone-EGFP and a heterozygous epithelial RFP reporter driven by the Krt14 promoter (Krt14p::RFP). Note that the surface-layer epithelial cells expressing high RFP comprise a cell sheet that folds inward during clefting.

**Video S5 (Related to Figure 2). 3D tracking of daughter cells from surface-derived cell divisions**. This video has 2 sequential parts. The first part shows “tracked cell highlighted” (left) and “tracked cell centered” (right) styles of an example cell tracking. The second part shows tracking of 16 pairs of daughter cells in the “tracked cell centered” style. Tracked cell is marked by a white dot. Distance indicated is from the coverslip. Time indicates minutes from the onset of anaphase (0 min).

**Video S6 (Related to Figure 6). Time-lapse videos of 3D spheroid cultures of engineered DLD-1 cells under indicated experimental conditions**. Images are merged phase contrast and epifluorescence channels. Green and magenta indicate cells expressing sg-Control or sg-Cdh1, respectively. Abscisic acid (ABA) is a dimerizer used to induce robust transcriptional repression in engineered DLD-1 cells. DMSO is the vehicle control. Live imaging started 5 days post Matrigel and ABA or DMSO treatment.

**Video S7 (Related to Figure 6). Time-lapse videos of a budding spheroid culture of engineered DLD-1 cells**. Images in the main window are maximum intensity projections of confocal fluorescence images, and images in the upper left inset are single-slice confocal fluorescence images. Green and magenta indicate cells expressing sg-Control or sg2-Cdh1, respectively. Yellow indicates the basement membrane marked by Atto-647N labeled fibronectin. Arrows and arrowheads indicate forming clefts. Live imaging started 4 days post Matrigel and ABA treatment.

## STAR METHODS

### Key Resources Table

| REAGENT or RESOURCE | SOURCE | IDENTIFIER |
| --- | --- | --- |
| **Antibodies** |  |  |
| Rat monoclonal anti-E-cadherin (clone ECCD-2) | Thermo Fisher | 13-1900 |
| Rat monoclonal anti-β1-integrin (clone mAb13) | MilliporeSigma | MABT821 |
| Rat monoclonal Anti-α5-integrin (clone mAb16) | MilliporeSigma | MABT820 |
| Goat polyclonal anti-collagen type IV | MilliporeSigma | AB769 |
| Rabbit polyclonal anti-laminin | MilliporeSigma | L9393 |
| Rabbit polyclonal anti-histone H3 (phospho S10) | Abcam | ab47297 |
| Rabbit monoclonal anti-β-catenin (clone E247) | Abcam | ab32572 |
| Rabbit Claudin 10 Polyclonal Antibody | Thermo Fisher | 38-8400 |
| Rabbit monoclonal anti-GAPDH (clone 14C10) | Cell Signaling | 2118L |
| Mouse monoclonal anti-α-tubulin (clone DM1α) | MilliporeSigma | T6199 |
| Rat Anti-Human CD49f α6-integrin (clone GoH3) | BD Biosciences | 555734 |
| Rat IgG2a, $\kappa$ Isotype Control (clone A110-2) | BD Biosciences | 553992 |
| Hamster Anti-Rat CD29 β1-integrin (clone Ha2/5) | BD Biosciences | 555002 |
| Hamster IgM, λ1 Isotype Control (clone G235-1) | BD Biosciences | 553957 |
| DyLight 405 donkey anti-rabbit | Jackson ImmunoResearch | 711-476-152 |
| Cy2 donkey anti-goat | Jackson ImmunoResearch | 705-225-147 |
| Alexa Fluor 488 donkey anti-rat | Jackson ImmunoResearch | 712-546-153 |
| Rhodamine Red-X donkey anti-rat | Jackson ImmunoResearch | 712-296-153 |
| Rhodamine Red-X donkey anti-rabbit | Jackson ImmunoResearch | 711-296-152 |
| Alexa Fluor 647 donkey anti-rat | Jackson ImmunoResearch | 712-606-153 |
| Alexa Fluor 647 donkey anti-rabbit | Jackson ImmunoResearch | 711-606-152 |
| 680RD goat anti-rabbit | LI-COR | 926-68071 |
| 800CW goat anti-rat | LI-COR | 926-32219 |
| 680RD goat anti-mouse | LI-COR | 926-68070 |
| 800CW goat anti-rabbit | LI-COR | 926-32211 |
| **Mouse Strains** |  |  |
| mTmG/HisG; FVB/N(Cg)-Gt(ROSA)26Sortm4(ACTB-tdTomato,-EGFP)Luo Tg(HIST1H2BB-EGFP)1Pa/Jhu | JHU (A. Ewald) | N/A |
| K14RFP/HisG; FVB/N(Cg)-Tg(KRT14-RFP)#Efu Tg(HIST1H2BB-EGFP)1Pa/Jhu | This study | N/A |
| KikGR; FVB/N(Cg)-Tg(KikGR) | Hsu et al., 2013 | N/A |
| ICR (CD-1) outbred mice | Envigo | Hsd:ICR(CD-1) |
| **Cell Lines** |  |  |
| 293T | Takara | 632273 |
| DLD-1 (parental) | ATCC | CCL-221 |
| Dia-C6 (DLD-1 expressing 4 transgenes for inducible transcriptional modulation) | This study | N/A |
| D188 (Dia-C6 with lenti-sg-Control and NLS-mScarlet) | This study | N/A |
| D193 (Dia-C6 with lenti-sg-Control and NLS-mNeonGreen) | This study | N/A |
| D266 (Dia-C6 with lenti-sg2-Cdh1 and NLS-mScarlet) | This study | N/A |
| D267 (Dia-C6 with lenti-sg1-Cdh1 and NLS-mScarlet) | This study | N/A |
| D301 (Dia-C6 with lenti-sg-Itgb1 and NLS-mNeonGreen) | This study | N/A |
| D2 (Dia-C6 with lenti-sg1-Cdh1, NLS-mScarlet, lenti-sg-Itgb1 and NLS-mNeonGreen) | This study | N/A |
| **Plasmids** |  |  |
| Super piggyBac Transposase expression vector | System Biosciences | PB210PA-1 |
| pSLQ2818 (PiggyBac-ABA-CRISPRi) | Addgene | 84241 |
| pW211 (pLenti-spsg-pEF1s-NLS-mNG-P2A-BlastR) | This study | N/A |
| pW212 (pLenti-spsg-pEF1s-NLS-mSL-P2A-BlastR) | This study | N/A |
| **Tissue Culture Media, Supplements and Enzymes** |  |  |
| DMEM | Thermo Fisher | 11965118 |
| RPMI-1640 | ATCC | 30-2001 |
| DMEM/F-12 | Thermo Fisher | 11039047 |
| Fetal Bovine Serum (FBS) | GE Healthcare / Cytiva | SH30070.03 |
| L-glutamine | Thermo Fisher | 25030081 |
| Penicillin-Streptomycin (PenStrep) | Thermo Fisher | 15140163 |
| L-Ascorbic Acid (Vitamin C) | MilliporeSigma | A7506 |
| Transferrin | MilliporeSigma | T8158 |
| Insulin-Transferrin-Selenium (ITS-G) (100×) | Thermo Fisher | 41400045 |
| Recombinant Mouse KGF/FGF-7 Protein | R&D Systems | 5028-KG-025 |
| Recombinant Mouse Neuregulin-1/NRG1 Protein | R&D Systems | 9875-NR-050 |
| Matrigel, Growth Factor Reduced Basement Membrane Matrix | Corning | 356231 |
| Laminin I, Cultrex 3D Culture Matrix (contains low amount of collagen IV) | Trevigen | 3446-005-01 |
| Trypsin-EDTA (0.05%), phenol red | Thermo Fisher | 25300120 |
| Accutase | Innovative Cell Technologies | AT104 |
| Dispase II | Thermo Fisher | 17105041 |
| Collagenase | Elastin Products Company | CL103 |

### Experimental Models and Method Details Mouse strains

All mouse experiments were performed under animal study protocols 14-745, 17-845, and 20-1040 approved by the NIDCR Animal Care and Use Committee (ACUC). All mouse embryos were used without sex identification (mixed sexes). All transgenic mice were in FVB/N background. The mT/mG;Histone-EGFP (Hadjantonakis and Papaioannou, 2004; Huebner et al., 2014; Muzumdar et al., 2007) mouse was a gift from A.J. Ewald (Johns Hopkins University). The Krt-14p::RFP (Zhang et al., 2011) mouse was a gift from M.P. Hoffman (NIDCR, NIH) and was originally from E. Fuchs (Rockefeller University). Krt-14p::RFP mice were crossed with mT/mG;Histone-EGFP mice to generate Krt-14p::RFP;Histone-EGFP mice. Our KikGR mouse was generated as described (Hsu et al., 2013). Transgenic mice 8-16 weeks old were bred to obtain 12- or 13-day old embryos. For timing of embryonic stage, the day after a vaginal plug was found was considered to be embryonic day 1. For experiments using wildtype embryos, timed pregnant ICR (CD-1) outbred mice were obtained from Envigo.

#### Cell lines

The HEK293T cell line used for lentivirus packaging was obtained from Takara (632273). The DLD-1 cell line was obtained from ATCC (CCL-221). DLD-1 cells were co-transfected with a PiggyBac transposase vector (System Biosciences, PB210PA-1) and a PiggyBac-ABA-CRISPRi vector (pSLQ2818; Addgene, 84241) using Lipofectamine 3000 (Thermo Fisher Scientific, L3000015), selected by 5 μg/mL puromycin (MilliporeSigma, P8833) and sorted for the presence of tagBFP to generate “Di” cells. Di cells were co-transfected with the above transposase vector, PiggyBac-ABA-CRISPRi vector and a modified PiggyBac-GA-CRISPRa vector (pW210, see **Plasmids**) using Lipofectamine 3000, selected by 5 μg/mL puromycin and 250 μg/mL hygromycin (MilliporeSigma, H3274) and sorted for brighter tagBFP than Di cells to generate “Dia” cells. Single cell clones of Dia cells were isolated by limiting dilution, and selected clones were functionally validated. The clonal Dia-C6 cells were used for lenti-sgRNA transduction followed by 20 μg/mL blasticidin (InvivoGen, ant-bl-1) selection and fluorescence cell sorting for mNeonGreen or mScarlet (see **Plasmids**) to obtain final DLD-1 derived cell lines used for spheroid culture experiments.

#### Plasmids

The modified PiggyBac-GA-CRISPRa vector (pW210) was generated by replacing the Zeocin resistance cassette of pSLQ2842 (Addgene, 84244) with a synthesized Hygromycin resistance cassette (IDT) by Gibson Assembly (Gibson et al., 2009). Lentiviral vectors for co-expressing sgRNAs and fluorescent nuclear reporters (pLenti-spsg-mNG/pW211, pLenti-spsg-mSL/pW212) were made by replacing the Cas9 expression cassette of lentiCRISPR v2 (Addgene, 52961) with an NLS-mNeonGreen-P2A-BlastR or NLS-mScarlet-I-BlastR cassette using Gibson Assembly. For lenti-sgRNA cloning, a pair of complementary oligos containing the desired sgRNA sequence (see **sgRNA design**) plus a 4-bp 5’-extension (“cacc” for the forward oligo and “aaac” for the reverse complementary oligo) was annealed to form an oligo duplex, which was ligated into Esp3I (NEB, R0734S) digested vectors by a 1:2 mixture of T4 ligase (NEB, M0202L) and T4 polynucleotide kinase (NEB, M0236L) in T4 ligase buffer. The ligation mix was transformed using NEB stable competent cells (NEB, C3040) for single colony isolation. The Miraprep (Pronobis et al., 2016) protocol was used to increase the yield of miniprep DNA, which was directly used for lentivirus packaging. Correct insertion of sgRNA sequence was confirmed by Sanger sequencing using primer 5’-gagggcctatttcccatgat-3’.

#### Salivary gland isolation and culture

Mouse submandibular salivary glands were isolated at embryonic day 12 or 13 (E12 or E13) as previously described (Sequeira et al., 2013). Briefly, a scalpel (Fine Science Tools, 10011-00 and 10003-12) was used to decapitate the mouse embryo. While the detached head was held on its side with one prong of forceps (Fine Science Tools, 11251-20) pierced through the top, a scalpel was used to slice across the mouth opening to isolate the mandible and tongue, between which the submandibular glands were sandwiched. Under a dissecting microscope, the detached mandible tissue was placed on a glass plate with the tongue facing down. A pair of forceps was used to slice through the midline of the mandible tissue to expose the tongue and the two submandibular glands attached to the base of the tongue. After surrounding tissues were removed, glands were detached using forceps and collected into a 35-mm dish with 3 mL DMEM/F-12 (Thermo Fisher, 11039047) media until all embryos were dissected. Isolated salivary glands were cultured on 13 mm diameter 0.1 μm pore polycarbonate filters (MilliporeSigma, WHA110405) floating on 200 μL Organ Culture Medium in the glass bottom area of a 50 mm MatTek dish (MatTek, P50G-1.5-14-F) at 37°C with 5% CO_2_. Organ Culture Medium was DMEM/F-12 supplemented with 150 μg/mL vitamin C (MilliporeSigma, A7506), 50 μg/mL transferrin (MilliporeSigma, T8158) and 1× PenStrep (100 units/mL penicillin, 100 μg/mL streptomycin; Thermo Fisher, 15140163).

#### Salivary gland collagenase treatment and washout

Paired salivary glands from the same embryo were separated into control and collagenase treatment groups. Purified collagenase (Elastin Products Company, CL103) was resuspended in water (Quality Biological, 351-029-131) for a 2 mg/mL stock (aliquoted and stored at -20°C). For collagenase washout, the polycarbonate filter with attached glands was transferred onto 2 mL fresh DMEM/F-12 in a 35 mm dish (Corning, 430165) and incubated for 15 min at 37°C for one wash. After 3× 15-min washes, the filter with glands was transferred onto 200 μL fresh Organ Culture Medium in a new 50 mm MatTek dish (see **Salivary gland isolation and culture**).

#### Isolation of salivary gland epithelial rudiments

Up to 6 intact salivary glands (see **Salivary gland isolation and culture**) were treated with 150 μL 2 units/mL dispase (Thermo Fisher, 17105041; diluted in DMEM/F-12) in a well of a Pyrex spot plate (Fisher Scientific 13-748B) for 15 min at 37°C. The glands were washed twice with 5% BSA (MilliporeSigma, A8577; diluted in DMEM/F-12) in the same well to quench the dispase activity. The mesenchyme of each gland was removed using a pair of forceps (Fine Science Tools, 11254-20) and a tungsten needle (Fine Science Tools, 10130-05 and 26016-12) under a dissecting microscope. The forceps were mostly used to hold the gland still, whereas the needle was gently inserted between the mesenchyme and epithelium to separate them. Isolated epithelial rudiments were transferred to a new well of the spot plate with 150 μL DMEM/F-12 media using low-retention pipette tips (cut for larger opening; Rainin, 30389190). When needed, a pair of forceps was used to cut off single epithelial buds from the isolated epithelial rudiments.

#### Single-cell dissociation of salivary gland epithelium

Single-cell dissociation of the salivary gland epithelium was performed as previously described with modifications (Sekiguchi and Hauser, 2019). Depending on experiments, 8-12 isolated epithelial rudiments from E13 submandibular salivary glands (see **Isolation of salivary gland epithelial rudiments and single epithelial buds**) were rinsed in 1 mL HBSS (Thermo Fisher, 14170161) in a 2 mL protein LoBind tube (Eppendorf, 022431102). After the rudiments were pelleted by centrifugation at 100× g for 30 seconds, liquid was removed as much as possible using a 1 mL pipette followed by a 200 μL pipette under a dissecting microscope to avoid accidentally discarding the samples. 100 μL Accutase (Innovative Cell Technologies, AT104) was added to the tube, which was immediately incubated for 2 minutes in a 37°C water bath to disrupt cell-cell adhesion. While being monitored under a dissecting microscope, the epithelial rudiments were triturated using a 200 μL pipet set at 50 μL using a low-retention tip (Rainin, 30389187) for 2 min, when most cells were clearly dissociated. 900 μL 10% fetal bovine serum (FBS; GE Healthcare/Cytiva, SH30070.03) diluted in PBS was added to quench the Accutase. The 1 mL cell suspension was passed through a 40 μm Flowmi (VWR, H13680-0040) cell strainer into a polyethylene terephthalate (PET) 15 mL tube (Corning, 430053). It is critical to use PET tubes instead of polypropylene tubes for efficient cell recovery. Cells were pelleted by centrifugation at 100× g for 2 minutes in a swinging-bucket rotor. After the supernatant was carefully removed, cells were resuspended in 1% FBS in PBS and pelleted again. About 50 μL liquid was retained to resuspend the cells, which was used for single cell capture (for scRNA-seq) or diluted in DMEM/F-12 media for 3D culture.

#### Single-cell RNA sequencing and data analysis

Single cell capture and library construction were performed using the 10x Genomics Chromium Single Cell 3’ Library & Gel Bead Kit (v2 Chemistry) following the manufacturer’s instructions. The libraries were sequenced on an Illumina NextSeq500 sequencer. The 10x Genomics Cell Ranger (v3.0.1) software suite was used for demultiplexing, read alignment and UMI (unique molecule identifier) counting. The filtered features, barcodes and matrix output files were used for further analysis using the Python libraries Scanpy (v1.6.0) (Wolf et al., 2018) and scVelo (v0.2.2) (Bergen et al., 2020). Scripts for cell clustering, cell cycle phase assignment and RNA velocity calculation are available on Github (see **DATA AND CODE AVAILABILITY**).

#### 3D culture of primary salivary gland epithelial buds or dissociated cells

Ultra-low attachment 96-well V-bottom (S-bio, MS-9096VZ) plates were used for 3D culture of salivary gland epithelial buds (see **Isolation of salivary gland epithelial rudiments**) or dissociated cells (see **Single-cell dissociation of salivary gland epithelium**). One epithelial bud or ∼3,000 dissociated epithelial cells were seeded per well in a 96-well plate with 50 μL DMEM/F-12 media. Immediately after seeding, the plate was centrifuged at 100×g for 3 min to sediment the epithelial bud or dissociated cells. 50 μL 2× culture mix containing 400 ng/mL FGF7 (R&D Systems, 5028-KG-025), 20 ng/mL NRG1 (R&D Systems, 9875-NR-050), 2× ITS-G supplement (Thermo Fisher, 41400045) and 1 mg/mL growth factor-reduced Matrigel (Corning, 356231; stock 9-10 mg/mL) were added to each well. Surrounding wells were filled with 100 μL HBSS to reduce culture media evaporation. The plate was cultured at 37°C with 5% CO_2_.

#### Single-molecule RNA fluorescence in situ hybridization (smFISH)

smFISH probe sets targeting Cdh1, Sox2 and Sox10 mRNAs were designed using the Stellaris probe designer and synthesized with either TAMRA-C9 or Quasar 670 dyes by LGC Biosearch Technologies. smFISH of wholemount E13 salivary glands was performed as previously described (Wang, 2018). Briefly, E13 salivary glands were fixed with 4% PFA in PBS at room temperature (RT) for 1 hour or overnight at 4°C, rinsed in PBSTx (PBS + 0.2% Triton-X-100), dehydrated sequentially in 30%, 50%, 70% and 100% methanol on ice, rehydrated sequentially in 70%, 50%, 30% methanol on ice, rinsed in PBSTx for 10 min at RT, permeabilized in 0.5% SDS in PBS at RT, equilibrated in smFISH Wash Solution (2× SSC and 10% formamide in DEPC-treated water) for 10 min at RT, hybridized in smFISH Hybridization Solution (2× SSC, 10% formamide, 10% dextran sulfate and 50 μg/mL yeast tRNAs in DEPC-treated water) containing 50 nM probes (1-2 nM each probe) at 37°C for 12 to 16 hours, washed in smFISH Wash Solution for 30 min at RT, stained with 0.5 μg/mL DAPI in smFISH Wash Solution for 2 hours at RT, washed 2 more times for 30 min at RT, rinsed in 2× SSC (30 mM sodium citrate and 300 mM sodium chloride; K D Medical, RGF-3240) and mounted in ProLong Diamond Anti-fade Mountant (Thermo Fisher, P36961) for imaging.

#### smFISH quantification

smFISH dot counting was performed using a suite of custom-written ImageJ macros as previously described (Wang, 2018). Briefly, smFISH images were smoothened by a Gaussian filter, contrast enhanced by a morphological top-hat filter (Legland et al., 2016), and local maxima points beyond a user-specified threshold level were identified and counted. An identical set of parameters was used to process all images from the same experiment.

#### 2D cell culture

DMEM (Thermo Fisher, 11965118) and RPMI-1640 (ATCC, 30-2001) media were supplemented with 10% fetal bovine serum (FBS; GE Healthcare/Cytiva, SH30070.03), 2 mM L-glutamine (Thermo Fisher, 25030081) and 1× PenStrep to make DMEM Complete and RPMI-1640 Complete media. Phenol red-free DMEM (GE Healthcare/Cytiva, SH30284.01) or RPMI-1640 (Thermo Fisher, 11835030) were used when cells were used for imaging or cell sorting. HEK293T cells were cultured in DMEM Complete medium in 37°C incubators with 10% CO_2_. DLD-1 and DLD-1 derived cells were cultured in RPMI-1640 Complete medium in 37°C incubators with 5% CO_2_. For passage, cells were detached using trypsin-EDTA (Thermo Fisher, 25300120) after rinsing with HBSS (Thermo Fisher, 14170161). Cell density was determined using an automated cell counter (Nexcelom Cellometer Auto 2000).

#### 3D spheroid culture

Ultra-low attachment 96-well U-bottom (Corning, 7007) or V-bottom (S-bio, MS-9096VZ) plates were used for 3D spheroid culture. DLD-1 cells expressing different sgRNAs were detached, pelleted by centrifugation at 1,000×g for 3 min, resuspended in RPMI-1640 Complete medium, counted and diluted to 60,000 cells/mL. For co-cultures of two cell types (e.g., sg-Control and sg-Cdh1 cells), appropriate volumes of the two cells were mixed in a separate tube to achieve desired mixing ratios. A multichannel pipette (Rainin, 17013810) was used to seed 50 μL cell suspensions in each well for 3,000 cells per spheroid. The 36 outer edge wells were filled with 100 μL HBSS to reduce medium evaporation over long culture periods (≥ 7 days). Immediately after seeding, the plate was centrifuged at 100×g for 3 min to sediment the cells. The next day, a 2× treatment mix of 44.8 μL RPMI-1640 Complete medium, 5 μL growth factor-reduced Matrigel (Corning, 356231; 9-10 mg/mL) and 0.2 μL DMSO (MilliporeSigma, D2650) or 50 mM abscisic acid (ABA; MilliporeSigma, A1049) was prepared for each well. A multichannel pipette was used to add 50 μL 2× treatment mix to each well for a final concentration of 5% Matrigel (450-500 μg/mL) and 100 μM ABA. Care was taken to minimize bubbles during pipetting. For integrin stimulation by MnCl_2_, 0.1 μL 50 mM MnCl_2_ was supplemented to every 50 μL 2× treatment mix. For integrin antibody blocking, two rat monoclonal antibodies (mAb13: anti-β1-integrin; mAb16: anti-α5-integrin) were diluted to 0.5 mg/mL in RPMI-1640 and passed through a desalting spin column (Thermo Fisher, 89883) pre-equilibrated for 4 times with RPMI-1640. Antibody concentrations were re-measured by absorbance at 280 nm on a nanodrop spectrophotometer (Denovix, DS-11). The per-well 2× treatment mix was adjusted to include 20 μL 0.5 mg/mL antibody solution, 19.4 μL RPMI-1640, 4.5 μL FBS, 0.45 μL 200 mM L-glutamine, 0.45 μL 100× PenStrep, 5 μL Matrigel and 0.2 μL 50 mM ABA.

#### sgRNA design

sgRNAs for target genes (Cdh1 or Itgb1) were designed on the CRISPOR (Haeussler et al., 2016) website using 500 bp sequences centered around the transcription start site (TSS ± 250 bp). sg1-Cdh1: 5’-gCCGAGAGGCTGCGGCTCCAA-3’. sg2-Cdh1: 5’-gTGGCCGGGGACGCCGAGCGA-3’. sg-Itgb1: 5’-GGACGCCGCGCGGAAAAGGT-3’. Control guide RNAs for both the S. pyogenes Cas9 (5’-gTGCGAATACGCCCACGCGAT-3’) and the S. aureus Cas9 (5’-gCCTTCCCAACAGTTGCGCAGC-3’) were designed from the bacterial lacZ gene against the human genome. An extra “g” was added to the 5’-end if the guide sequence did not begin with “g” to facilitate transcription by the U6 promoter.

#### Lentivirus packaging

All lentivirus work was performed using in a BSL2 room with a dedicated incubator. Lenti-sgRNA vectors (see **Plasmids**) were co-transfected with psPAX2 (Addgene, 12260) and pMD2.G (Addgene, 12259) into HEK293T cells by calcium co-precipitation to produce infectious lentiviral particles. Briefly, 4×10^6^ HEK293T cells were seeded in a 10 cm dish one day before packaging. Next morning, culture media were changed and supplemented with 25 μM chloroquine (MilliporeSigma, C6628). Two 15 mL tubes (A and B) were used to prepare the transfection mix. 1 mL 2× HBS (50 mM HEPES, 280 mM NaCl, 1.5 mM Na_2_HPO_4_, pH 7.10) was added to tube A. 10 μg of each plasmid (the lenti-sgRNA vector, psPAX2 and pMD2.G) and 1 mL 0.3 M CaCl_2_ were sequentially added to tube B and mixed by pipetting. The DNA-CaCl_2_ mixture in tube B was then added dropwise into the 2× HBS in tube A and mixed by pipetting. The transfection mix was then added dropwise to the 10 cm dish. Culture media were changed twice at about 12- and 36-hours post transfection, and lentivirus-containing media were collected twice at about 36- and 60-hours post transfection into a 50 mL tube (stored at 4°C). Pooled lentivirus-containing media were passed through a 0.45 μm filter (MilliporeSigma, SE1M003M00) to remove cell debris. To concentrate the lentivirus, 4 mL 5× PEG reagent (System Biosciences, LV825A-1) was added and mixed by pipetting. After ≥ 12 hours incubation at 4°C (up to 4 days), lentivirus was pelleted by centrifugation at 1,500× g for 30 min at 4°C, and the pellet was resuspended in 400 μL DMEM/F-12 with 1× PenStrep and stored at -80°C.

#### Lentivirus titration

The titer of concentrated lentivirus was estimated using Lenti-X GoStix Plus (Takara, 631281) after 100× dilution. A GoStix Value (GV) of 50 was empirically considered to be equivalent to a lentivirus titer of 5×10^5^ IFU/mL. The typical titer of concentrated sgRNA lentivirus was 1.5×10^8^ IFU/mL.

#### Lentivirus transduction

One day before transduction, 1×10^5^ cells were seeded in a well of a 12-well plate. Next day, 2 μL 4 mg/mL polybrene (MilliporeSigma, H9268) was added to 1 mL medium (final 8 μg/mL). An appropriate amount of lentivirus for an MOI (multiplicity of infection; ratio of infectious viral particles to cells) of 10-15 (typically 20 μL for concentrated sgRNA lentivirus) was then added. One day later, the virus-containing medium was replaced with regular medium after 4× HBSS washes, and cells were re-plated to a 75 cm^2^ flask in culture medium supplemented with 20 μg/mL blasticidin (InvivoGen, ant-bl-1) to begin the antibiotic selection.

#### Fluorescence activated cell sorting

For cell sorting, DLD-1 derived cells were trypsinized for 5 min longer than for passage (∼15 min total) to increase the ratio of single cells, pelleted at 1,000× g for 3 min, and resuspended in phenol-red free, serum-free RPMI-1640 medium (Thermo Fisher, 11835030) for a cell density of 5-10×10^6^/mL. The cell suspension was passed through a 40 μm Flowmi cell strainer (VWR, H13680-0040) and sorted on a BD FACSAria III or SONY SH800 cell sorter operated by the NIDCR Combined Technical Research Core.

#### Western blotting

DLD-1 derived cells were seeded in 12-well plates (Corning, 3512) at 20,000 cells/well on day 0, treated with DMSO vehicle or desired concentrations of ABA in DMSO on day 1 (or each day of days 1-5 per well for an ABA time course), and harvested on day 6. Culture media were changed once on day 4 or 5. For harvesting, 100 μL RIPA buffer (25 mM Tris, pH 7.4, 150 mM NaCl, 1% NP-40, 0.5% sodium deoxycholate, 0.1% SDS) supplemented with protease inhibitors (MilliporeSigma, 11836170001) was added to each well after rinsing with PBS (Phosphate Buffered Saline; Lonza, 17-517Q). Cells were scraped into RIPA buffer on ice using 1 mL pipette tips. Cell suspensions were transferred to pre-cooled 1.5 mL tubes (Eppendorf, 022363212), incubated for 30 min on ice, and centrifuged at 13,000× rpm for 10 min at 4°C. Cleared cell lysates were transferred to a new set of pre-cooled 1.5 mL tubes and stored at -20°C. Protein concentrations of cell lysates were determined by Bradford assays (Bio-Rad, 5000201). Lysate aliquots with 16 μg protein were denatured in 1× Laemmli sample buffer (Bio-Rad, 1610747) for 5 min at 95°C. Samples of 8 μg protein or 10 μL protein ladder (Thermo Fisher, 26623) were loaded per lane onto a precast gel (Bio-Rad, 4561096) for electrophoresis. Proteins were transferred onto a nitrocellulose membrane (Bio-Rad, 1704159) using the Turbo Transfer system (Bio-Rad, 1704150). The membrane was stained with Ponceau S (MilliporeSigma, P7170) to assess transfer quality, washed for 5 min in TBST (Tris Buffered Saline with 0.1% Tween-20; Quality Biological, 351-086-101; MilliporeSigma, P2287) to remove Ponceau staining, blocked in Blocking Solution (5% nonfat dry milk in TBST) for 30 min at room temperature (RT), incubated in primary antibodies (see **Antibody usage for Western blotting**) diluted in Blocking Solution overnight at 4°C, washed 4× 5 min with TBST at RT, incubated in LI-COR secondary antibodies diluted in Blocking Solution for 1-2 hours at RT, washed 4× 5 min in TBST at RT, and imaged on a LI-COR Odyssey CLx imaging system controlled by LI-COR Image Studio software. Western blotting band intensities were quantified using LI-COR Image Studio Lite software.

#### Antibody usage for Western blotting

Primary antibodies used for Western blotting: anti-E-cadherin (Thermo Fisher, 13-1900), 0.5 μg/mL; anti-β-catenin (Abcam, ab32572), 1:5,000 (0.0126 μg/mL); anti-β1-integrin (MilliporeSigma, MABT821), 1 μg/mL; anti-GAPDH (Cell Signaling, 2118L), 1:2,000; anti-α-tubulin (MilliporeSigma, T6199), 0.5 μg/mL. All Western blotting secondary antibodies were from LI-COR and used at 1:5,000 for 800CW conjugates and 1:10,000 for 680RD conjugates.

#### Immunostaining of cells

Cells were seeded, immunostained and imaged in 8-well ibidi chambers (ibidi, 80826). All procedures were performed at room temperature with gentle rocking. Cells were fixed with 4% PFA in PBS for 15 min, permeabilized with PBSTx for 15 min, blocked in 5% donkey serum in PBS for 30 min, incubated in primary antibodies (see **Antibody usage for immunostaining**) diluted in PBS for 1 hour, washed 4× 5 min with PBS, incubated in 0.5 μg/mL DAPI (Thermo Fisher, D1306) and secondary antibodies diluted in PBS for 1 hour, washed 4× 5 min with PBS, stored at 4°C and imaged within 3 days.

#### Immunostaining of spheroids and salivary glands

Spheroids were rinsed in 2 mL PBS in a 35 mm dish and transferred into sample baskets (one basket per staining group; Intavis, 12.440) using low-retention pipette tips (cut for larger opening; Rainin, 30389187 or 30389190) under a dissecting microscope. For fixation, each basket was soaked in 1 mL fixative (4% PFA in PBS; Electron Microscopy Sciences, 15710) in a well of a 24-well plate (Corning, 3524) overnight at 4°C. Cultured salivary glands were fixed on the filter by replacing Organ Culture Medium under the filter with 200 μL fixative for 1 hour at room temperature (RT) or overnight at 4°C. Fixed glands were detached from the filter and transferred into sample baskets in PBS in a 35 mm dish using a pair of forceps (Fine Science Tools, 11251-20) under a dissecting microscope. Fixed samples in baskets were permeabilized in PBSTx (PBS with 0.2% Triton-X-100; Thermo Fisher, 28314) for 30 min at RT, blocked in 5% donkey serum (Jackson ImmunoResearch, 017-000-121) in PBSTx for 2 hours at RT, incubated in primary antibodies (see **Antibody usage for immunostaining**) diluted in either PBSTx or 5% donkey serum for 2 days at 4°C, washed 4× 15 min in PBSTx at RT, incubated in secondary antibodies diluted in either PBSTx or 5% donkey serum for 2 days at 4°C, washed 4× 15 min in PBSTx at RT, rinsed in PBS and mounted under a dissecting microscope. To preserve fluorescence and to minimize compression, samples were mounted in 20 or 40 μL antifade mountant (Thermo Fisher, P36930) supported by one layer (for salivary glands) or two layers (for spheroids) of imaging spacers (Grace Bio-labs, 654004) attached to a glass slide (Thermo Fisher, 3011-002).

#### Antibody usage for immunostaining

Primary antibodies used for immunostaining: anti-E-cadherin (Thermo Fisher, 13-1900), 1 μg/mL; anti-collagen type IV (MilliporeSigma, AB769), 2 μg/mL; anti-laminin (MilliporeSigma, L9393), 2.5 μg/mL; anti-histone H3 (phospho S10) (Abcam, ab47297), 1 μg/mL; anti-β1-integrin (MilliporeSigma, MABT821), 1 μg/mL; anti-Claudin 10 (Thermo Fisher, 38-8400), 1 μg/mL. All immunostaining secondary antibodies were from Jackson ImmunoResearch (an equal volume of glycerol was added for storage at -20°C after reconstitution as instructed) and used at 1:200 (1.5-3 μg/mL).

#### Integrin blocking for salivary gland culture

Hamster anti-β1-integrin (BD Biosciences,), isotype-matched control IgM (BD Biosciences,), rat anti-α6-integrin (BD Biosciences, 555734) and isotype-matched control IgG (BD Biosciences, 553992) were used at 100 μg/mL each for function blocking or control after overnight dialysis into 1 liter DMEM/F-12 media supplemented with 1× PenStrep at 4°C with gentle agitation, using a dialysis cassette (Thermo Fisher, 66383).

#### Integrin blocking for 3D DLD-1 spheroid culture

Rat anti-β1-integrin (in-house clone mAb13; available from MilliporeSigma, MABT821) was used at 100 μg/mL for function blocking. Rat anti-α5-integrin (in-house clone mAb16; available from MilliporeSigma, MABT820) was used at 100 μg/mL as a control in the β1-integrin function blocking experiment. Stock antibodies (1.9-5 mg/mL) were diluted to 0.5 mg/mL in RPMI-1640 media and further exchanged to RPMI-1640 media using spin desalting columns (Thermo Fisher, 89882) following the manufacturer’s instructions.

#### Tissue clearing of spheroids

Tissue clearing was performed for the images shown in **Fig. 6G** to enable imaging over 200 μm thickness. For tissue clearing, regular immunostaining steps were carried out except for mounting. Spheroids were instead sequentially transferred to each well of a 3-well silicone chamber slide (ibidi, 80381), each containing 500 μL CytoVista clearing reagent (Thermo Fisher, V11315). After 5 min incubation in the last well, spheroids were transferred to 200 μL CytoVista clearing reagent per well of an 8-well glass-bottom ibidi chamber (ibidi, 80827) for imaging with two-photon microscopy (see **Immunostaining light microscopy**). Note that this clearing reagent preserved mScarlet but not mNeonGreen fluorescence.

#### Cell attachment assay

DLD-1 derived cells (sg-Control, sg1-Cdh1, sg2-Cdh1 and sg-Itgb1) were pre-treated with 100 μM ABA for 5-10 days before being used. The glass surface of MatTek 6-well plates (MatTek, P06G-0-14-F) was coated with 200 μL 91 μg/mL Matrigel in PBS or 8 μg/mL E-cadherin extracellular domain (E-cad-ECD; R&D Systems, 8505-EC-050) in PBS for 1-3 hours at 37°C. Coated wells were rinsed once with 3 mL PBS, blocked with 2 mL 5% bovine serum albumin (BSA; MilliporeSigma, 10735108001) in PBS for 30 min at 37°C, and washed twice with 3 mL PBS. During the blocking step, cells were detached, pelleted, resuspended, counted and diluted to 5×10^4^ cells/mL. 2 mL cell suspension was seeded in each well. 3 or 4 wells were used per experimental group. After 15 min (for Matrigel coating) or 2 hours (for E-cad-ECD coating) incubation at 37°C with 5% CO_2_, unattached cells were removed from each well, which was then gently washed 3× with 3 mL PBS, fixed with 2 mL 4% PFA in PBS for 15 min at 37°C, washed 2× with 3 mL PBS, and imaged under a microscope (see **Live-spheroid imaging and cell-attachment assay imaging**) to quantify attached cell densities (see **Image processing and analysis**). For assay consistency, a 2 mL aspirating pipette was capped with a 200 μL pipette tip to attenuate vacuum strength, and the house vacuum valve was pre-adjusted using PBS to reach a liquid removal rate of ∼1 mL/second. During liquid removal and addition, tips of aspirating or transferring pipettes were always placed to the side of the bottom of the MatTek well away from the coated glass surface, resulting in ∼200 μL leftover liquid between washes. Care was taken throughout the assay to avoid agitating the plates. The incubation time for each coating was determined to be the time at which ∼50% of control cells were attached in pilot assays.

#### Live-organ imaging by two-photon microscopy

All microscopy systems for live imaging were equipped with an environmental chamber to maintain samples at 37°C with 50% humidity and 5% CO_2_. Transgenic salivary glands expressing fluorescent markers were isolated and cultured on a floating 13 mm filter at 37°C with 5% CO_2_ (see **Salivary gland isolation and culture**) for at least 1 hour before being mounted for live imaging. Double-adhesive imaging spacers (Grace Bio-labs, 654008; cut into 8 separated wells, each 120 μm thick with 9 mm diameter opening) were sterilized by soaking in 70% ethanol for 3 min and attached to the glass bottoms of 50 mm MatTek dishes (MatTek, P50G-1.5-30-F). Under a dissecting microscope, 5 μL Organ Culture Medium was transferred to the center of the imaging spacer, and the filter with glands was flipped onto the imaging spacer so that glands were sandwiched between the filter and the glass bottom. Care was taken to ensure the filter was flat and center-aligned with the imaging spacer. The edge of the filter was pressed to ensure tight adherence to the imaging spacer. 2 mL Organ Culture Medium was then added to the MatTek dish, which was incubated at 37°C with 5% CO_2_ for at least 2 hours before imaging. A Nikon 40×, 1.15 NA, Apo LWD, water-immersion objective or a Nikon 25×, 1.05 NA, Plan Apo, silicone-immersion objective was used for live-organ imaging using two-photon microscopy on a Nikon A1R Confocal Microscope System equipped with a Ti:sapphire laser (Coherent, Chameleon Vision II). Image acquisition was controlled by Nikon NIS-Elements software. Images were acquired at 2 μm z intervals over 100 μm thickness and 5 min intervals for 20-36 hours. The tunable laser was used at 950 nm for simultaneous two-photon excitation of histone-EGFP and membrane-tdTomato (or heterozygous Krt14p::RFP), and 920 nm for histone-EGFP and homozygous Krt14p::RFP. The laser power was adjusted to compensate for z-depth changes using the “Z Intensity Correction” option with criteria that the bulk histograms of both channels spanned 500-1500 gray values (at typically 1-8% or 12-100 mW power).

#### KikGR photoconversion and live imaging by confocal microscopy

E13 KikGR transgenic glands were cultured on the filter with Organ Culture Medium supplemented with 5 μg/mL AF680-anti-collagen IV (see **Protein labeling**) for ∼10 hours. Under a dissecting microscope, a 10 μL pipette tip was used to spot vacuum grease (MilliporeSigma, 18405) around glands on the filter, which was then flipped onto the glass area of a 35 mm dish (MatTek, P35G-1.5-20-C). The vacuum grease limited gland compression and also served as a bio-inert glue to adhere the filter to the glass. 2 mL Organ Culture Medium supplemented with 1 μg/mL AF680-anti-collagen IV (see **Protein labeling**) were added to the dish for imaging after 2 hours incubation at 37°C with 5% CO_2_. A Nikon 40×, 1.15 NA, Apo LWD, water-immersion objective was used for photoconversion and imaging on a Nikon A1R Confocal Microscope System equipped with 4 laser lines (405 nm, 488 nm, 561 nm, 640 nm; Nikon LU-N4). Photoconversion and image acquisition were controlled by Nikon NIS-Elements software. For photoconversion, a 405 nm laser was used at 1-5% power (0.15-0.75 mW) with the pinhole set at 1 AU (Airy Unit; 30.7 μm) to stimulate user-specified polygonal ROIs (Regions of Interest) in the “ND Stimulation” module of the software. Short 1-5 second pulses (depending on ROI sizes) were repeated until all green fluorescence inside ROIs was converted to red fluorescence. For image acquisition, 488 nm and 561 nm lasers were used at 2% power (0.3 mW), whereas the 640 nm laser was used at 5% (0.75 mW) with the pinhole set at 1.2 AU (58.7 μm). Images were acquired at 10 min time intervals and 2 μm z intervals.

#### Live-spheroid imaging and cell-attachment assay imaging

Live-spheroid imaging (time-lapse or single-time-point) and cell-attachment assay imaging were performed by phase contrast and epifluorescence microscopy using a Nikon 10×, 0.3 NA, Plan Fluor objective on a Nikon Ti-E brightfield microscope system with a Hamamatsu Orca Flash 4.0 V3 sCMOS camera. Spheroids were imaged in the same 96-well U-bottom or V-bottom plates for culture. Image acquisition was controlled by Nikon NIS-Elements software. The JOBS module of the software was used to automatically set up multiple positions in a 96-well plate (spheroids) or a 6-well plate (cell attachment assay). For time-lapse live-spheroid imaging by confocal microscopy, spheroids were incubated in 40 μg/mL Atto647N-fibronectin (see **Protein labeling**) overnight at 37°C with 5% CO_2_ and transferred into a 4-well 35 mm dish (ibidi, 80466), each well of which contained 2 or 3 spheroids in 100 μL spheroid culture medium with 10 μg/mL Atto647N-fibronectin. A Nikon 20×, 0.75 NA, Plan Apo objective was used on a Nikon A1R Confocal Microscope System equipped with 4 laser lines. A resonant scanner was used for high-speed laser scanning. Image acquisition was controlled by Nikon NIS-Elements software at 10 min time intervals and 2 μm z intervals.

#### Immunostaining light microscopy

Immunostained spheroids in **Fig. 6G** were imaged by two-photon microscopy using a Nikon 20×, 0.75 NA objective on a Nikon A1R Confocal Microscope System equipped with a Ti:sapphire laser (used at 760 nm, 3% or 90 mW) controlled by Nikon NIS-Elements software. All other immunostained spheroids and salivary glands were imaged by laser scanning confocal microscopy using Nikon 20×, 0.75 NA or 60×, 1.4 NA Plan Apo objectives on a Nikon A1R Confocal Microscope System controlled by Nikon NIS-Elements software or a Zeiss 20×, 0.75 NA or 63×, 1.4 NA Plan Apo objective on a Zeiss LSM 880 system controlled by Zeiss ZEN software. Immunostained tissue culture cells were imaged using a Nikon 40×, 1.25 NA, Plan Apo, silicone-immersion objective on a Nikon spinning disk confocal system equipped with a Yokogawa CSU-X1 unit and a Prime 95B sCMOS camera (Photometrics) controlled by Nikon NIS-Elements software.

#### Protein labeling

Human plasma fibronectin was purified as previously described (Akiyama, 1999). For fibronectin labeling, NHS-Atto647N (ATTO-TEC, AD 647N-31) was dissolved in DMSO to 10 mM and stored in a desiccated container at -20°C as 10 μL aliquots. Labeling buffer was 20 parts PBS with 1 part 0.2 M NaHCO_3_ (pH 9.0), which was adjusted to pH 8.3. For labeling, 1 mL fibronectin was exchanged to the labeling buffer using a spin desalting column (Thermo Fisher, 89891). The fibronectin concentration after buffer exchange was determined by absorbance at 280 nm (2.3 mg/mL), which was used to calculate the amount of NHS-Atto647N equivalent to 4× times molar excess of fibronectin (3.4 μL). The labeling mix was incubated for 45 min at room temperature, then unlabeled dye was removed by buffer exchange into PBS using a spin desalting column. Insoluble Atto647N-fibronectin was removed by centrifugation at 13,000× rpm for 30 min at 4°C. The concentration of cleared Atto647N-fibronectin and the degree of labeling (DOL) were calculated from absorbance at 280 nm and 646 nm with a dye-specific correction factor 0.03 (using the Atto calculation formula; 0.84 mg/mL, DOL 1.7). Atto647N-fibronectin was then aliquoted and stored at -80°C. A similar procedure was used to label collagen IV antibody (MilliporeSigma, AB769) with NHS-AF680 (Thermo Fisher, A20008), but a 50 mM borate buffer (pH 9.0) was used as the labeling buffer and labeled antibodies were stored in PBS at 4°C.

#### Image processing and analysis

All images acquired on the Nikon A1R Confocal System were denoised using the Denoise.ai function of the Nikon NIS-Elements software. All other image processing was performed in Fiji (Schindelin et al., 2012), an ImageJ distribution. Customized Python, Jython and ImageJ Macro scripts were used for automating or facilitating image analysis and data visualization in this study (see **DATA AND CODE AVAILABILITY**). Automatic 3D cell tracking and surface rendering were performed using Imaris 9.5.0 (Bitplane). Manual curation of semi-automatic cell tracking was performed in Fiji using the TrackMate (Tinevez et al., 2017) plugin. Manual surface reconstruction was performed by drawing polylines along the epithelial surface at sparse z planes (∼6 μm intervals) on x-y view and resliced y-z view image stacks in Fiji, which were used for interpolation and plotting using customized Python scripts. For automated cell counting in the attachment assay, nuclear fluorescence images were smoothed by Gaussian filter (sigma = 2 pixels), contrast enhanced by the MorphoLibJ (Legland et al., 2016) white top hat filter (disk, radius = 10), binarized and counted using the Analyze Particles function of ImageJ. For manual bud counting, several efforts were made to minimize bias. First, file names of all images were scrambled before counting for observer blinding. Second, the same investigator performed the counting of all spheroids (K.M.) or all salivary glands (S.W.) to avoid between-person variance. Third, an explicit criterion was used such that a bud was counted only when its protruding edge occupied at least one third of a circle.

